# Spectrum preserving tilings enable sparse and modular reference indexing

**DOI:** 10.1101/2022.10.27.513881

**Authors:** Jason Fan, Jamshed Khan, Giulio Ermanno Pibiri, Rob Patro

## Abstract

The reference indexing problem for *k*-mers is to pre-process a collection of reference genomic sequences *ℛ* so that the position of all occurrences of any queried *k*-mer can be rapidly identified. An efficient and scalable solution to this problem is fundamental for many tasks in bioinformatics.

In this work, we introduce the *spectrum preserving tiling* (SPT), a general representation of *ℛ* that specifies how a set of *tiles* repeatedly occur to `spell` out the constituent reference sequences in *ℛ*. By encoding the order and positions where *tiles* occur, SPTs enable the implementation and analysis of a general class of modular indexes. An index over an SPT decomposes the reference indexing problem for *k*-mers into: (1) a *k*-mer-to-tile mapping; and (2) a tile-to-occurrence mapping. Recently introduced work to construct and compactly index *k*-mer sets can be used to efficiently implement the *k*-mer-to-tile mapping. However, implementing the tile-to-occurrence mapping remains prohibitively costly in terms of space. As reference collections become large, the space requirements of the tile-to-occurrence mapping dominates that of the *k*-mer-to-tile mapping since the former depends on the amount of total sequence while the latter depends on the number of unique *k*-mers in *ℛ*.

To address this, we introduce a class of sampling schemes for SPTs that trade off speed to reduce the size of the tile-to-reference mapping. We implement a practical index with these sampling schemes in the tool `pufferfish2`. When indexing over 30,000 bacterial genomes, `pufferfish2` reduces the size of the tile-to-occurrence mapping from 86.3GB to 34.6GB while incurring only a 3.6× slowdown when querying *k*-mers from a sequenced readset.

**Supplementary materials:** Sections S.1 to S.8 available online at https://doi.org/10.5281/zenodo.7504717

**Availability:** `pufferfish2` is implemented in Rust and available at https://github.com/COMBINE-lab/pufferfish2.

## 1 Introduction

Indexing of genomic sequences is an important problem in modern computational genomics, as it enables the atomic queries required for analysis of sequencing data — particularly *reference guided* analyses where observed sequencing data is compared to known *reference* sequences. Fundamentally, analyses need to first rapidly locate short exact matches to reference sequences before performing other operations downstream. For example, for guided assembly of genomes, variant calling, and structural variant identification, seed sequences are matched to known references before novel sequences are arranged according to the seeds [1]. For RNA-seq, statistics for groups of related *k*-mers mapping to known transcripts or genes allow algorithms to infer the activity of genes in single-cell and bulk gene-expression analyses [2,3,4].

Recently, researchers have been interested in indexing collections of genomes for metagenomic and pan-genomic analyses. There have been two main types of approaches: full-text indexes, and hashing based approaches that typically index the *de Bruijn graph* (dBG). With respect to full-text indexes, researchers have developed tools that use the *r-index* [5] to compute matching statistics and locate maximal exact matches for large reference collections [6,7]. For highly repetitive collections, such as many genomes from the same species, r-index based approaches are especially space efficient since they scale linearly to the number of runs in the *Burrows-Wheeler Transform* (BWT) [8] and not the length of the reference text. With respect to hashing based approaches, tools restrict queries to fixed length *k*-mers [1,9] and index the dBG. These tools achieve faster exact queries but typically trade off space. In other related work, graph-based indexes that compactly represent genomic variations as paths on graphs have also been developed [10,11]. However, these indexes require additional work to project queries landing on graph-based coordinates to linear coordinates on reference sequences.

Many tools have been developed to efficiently build and represent the dBG [12,13]. Recently, Khan et al. introduced a pair of methods to construct the compacted dBG from both assembled references [14] and read sets [15]. Ekim et al. [16] introduced the minimizer-space dBG — a highly effective lossy compression scheme that uses minimizers as representative sequences for nodes in the dBG. Karasikov et al. developed the Counting dBG [17] that stores differences between adjacent nodes in the dBG to compress metadata associated with nodes (and sequences) in a dBG. Encouragingly, much recent work on *Spectrum Preserving String Sets* (SPSS) that compactly index the set-membership of *k*-mers in reference texts has been introduced [18,19,20,15,21,22,23]. Although these approaches do not tackle the *locate* queries directly, they do suggest that even more efficient solutions for reference indexing are possible.

In this work, we extend these recent ideas and introduce the concept of a *Spectrum Preserving Tiling* (SPT) which encodes how and where *k*-mers in an SPSS occur in a reference text. In introducing the SPT, this work makes two key observations. First, a hashing based solution to the reference indexing problem for *k*-mers does not necessitate a de Bruijn graph but instead requires a *tiling* over the input reference collection — the SPT formalizes this. Second, the reference indexing problem for *k*-mers queries can be cleanly decomposed into a *k-mer-to-tile* query and a *tile-to-occurrence* query. Crucially, SPTs enable the implementation and analysis of a general class of modular indexes that can exploit efficient implementations introduced in prior work.

### Contributions

We focus our work on considering how indexes can, *in practice*, efficiently support the two composable queries — the *k-mer-to-tile* query and the *tile-to-occurrence* query. We highlight this work’s key contributions below. We introduce:

1. The *spectrum preserving tiling* (SPT). An SPT is a general representation that explicitly encodes how shared sequences — *tiles* — repeatedly occur in a reference collection. The SPT enables an entire *class* of sparse and modular indexes that support exact locate queries for *k*-mers.
2. An algorithm for sampling and compressing an indexed SPT built from unitigs that *samples* unitig-occurrences. For some small constant “sampling rate”, *s*, our algorithm stores the positions of only ≈ 1/*s* occurrences and encodes all remaining occurrences using a small *constant* number of bits.
3. `Pufferfish2:` a practical index and implementation of the introduced sampling scheme. We highlight the critical engineering considerations that make `pufferfish2` effective in practice.

## 2 Problem definition and preliminaries

### The mapped reference position (MRP) query

In this work we consider the *reference indexing problem for k-mers*. Given a collection of references *ℛ* = {*R*_1_, …, *R*_*N*_ }, where each reference is a string over the DNA alphabet `{A, C, T, G}`, we seek an index that can efficiently compute the *mapped reference position* (MRP) query for a fixed *k*-mer size *k*. Given any *k*-mer *x*, the MRP query enumerates the positions of all occurrences of *x* in *ℛ*. Precisely, each returned occurrence is a tuple (*n, p*) that specifies that *k*-mer, *x*, occurs in reference *n* at position *p* where *R*_*n*_[*p* : *p* + *k*] = *x*. If a *k*-mer does not occur in some *R*_*n*_ ∈ *ℛ*, the MRP query returns an empty list.

### Basic notation

Strings and lists are zero-indexed. The length of a sequence *𝒮* is denoted |*𝒮*|. The *i*-th character of a string *𝒮* is *𝒮*[*i*]. A *k*-mer is a string of length *k*. A sub-string of length *ℓ* in the string *𝒮* starting at position *i* is notated *𝒮*[*i* : *i* + *ℓ*]. The prefix and suffix of length *i* is denoted *𝒮*[: *i*] and *𝒮*[|*𝒮*| − *i* :], respectively. The concatenation of strings *A* and *B* is denoted *A* ∘ *B*.

We define the *glue* operation, *A* ⊕_*k*_ *B*, to be valid for any pair of strings *A* and *B* that overlap by (*k* − 1) characters. If the (*k* − 1)-length suffix of *A* is equal to the (*k* − 1)-length prefix of *B*, then *A* ⊕_*k*_ *B* := *A* ∘ *B*[(*k* − 1) :]. When *k* clear from context, we write *A* ⊕ *B* in place of *A* ⊕_*k*_ *B*.

### Rank and select queries over sequences

Given a sequence *𝒮*, the *rank* query given a character *α* and position *i*, written `rank`_*a*_(*𝒮, i*), is the number of occurrences of *α* in *𝒮*[: *i*] The *select* query `select`_*a*_(*𝒮, r*) returns the position of the *r*-th occurrence of symbol *α* in *𝒮*. The *access* query `access`(*𝒮, i*) returns *𝒮*[*i*]. For a sequence of length *n* over an alphabet of size *σ*, these can be computed in *𝒪*(lg *σ*) time using a *wavelet matrix* that requires *n* lg *σ* + *o*(*n* lg *σ*) bits [24].

## 3 Spectrum preserving tilings

In this section, we introduce the *spectrum preserving tiling*, a representation of a given reference collection *ℛ* that specifies how a set of *tiles* containing *k*-mers repeatedly occur to spell out the constituent reference sequences in *ℛ*. This alternative representation enables a modular solution to the reference indexing problem, based on the interplay between two mappings — a *k*-mer-to-tile mapping and a tile-to-occurrence mapping.

### 3.1 Definition

Given a *k*-mer length *k* and an input reference collection of genomic sequences *ℛ* = {*R*_1_, …, *R*_*N*_ }, a spectrum preserving tiling (SPT) for *ℛ* is a five-tuple Γ ? (*𝒰, 𝒯, 𝒮, 𝒲, 𝔏*):

- **Tiles**: *𝒰* = {*U*_1_, …, *U*_*F*_ }. The set of *tiles* is a spectrum preserving string set, i.e., a set of strings such that each *k*-mer in *ℛ* occurs in some *U*_*i*_ ∈ *ℛ*. Each string *U*_*i*_ ∈ *𝒰* is called a *tile*.
- **Tiling sequences**: *𝒯* = {*T*_1_, …, *T*_*N*_ } where each *T*_*n*_ corresponds to each reference *R*_*n*_ ∈ *ℛ*. Each tiling sequence is an ordered sequence of tiles 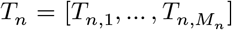 of length *M*_*n*_, with each *T*_*n,m*_ = *U*_*i*_ ∈ *𝒰*. We term each *T*_*n,m*_ a *tile-occurrence*.
- **Tile-occurrence lengths**: *𝔏* = {*L*_1_, …, *L*_*N*_ }, where each 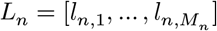is a sequence of lengths.
- **Tile-occurrence offsets**: *𝒲* = {*W*_1_, …, *W*_*N*_ }, where each 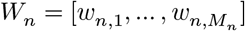] is an integer-sequence.
- **Tile-occurrence start positions**: *𝒮* = {*𝒮*_1_, …, *𝒮*_*N*_ }, where each 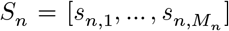 is an integer-sequence.

A valid SPT must satisfy the *spectrum preserving tiling property*, that every reference sequence *R*_*n*_ can be reconstructed by gluing together *substrings of tiles* at offsets *W*_*n*_ with lengths *L*_*n*_:

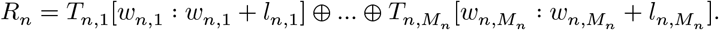

Specifically, the SPT encodes how redundant sequences — *tiles* — repeatedly occur in the reference collection *ℛ*. We illustrate how an ordered sequence of start-positions, offsets, and lengths explicitly specify how redundant sequences tile a pair of references in Fig. 1. More succinctly, each tile-occurrence *T*_*n,m*_ with length *l*_*n,m*_ tiles the reference sequence *R*_*n*_ as:

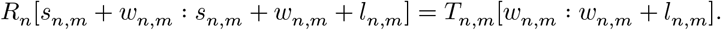

In the same way a small SPSS compactly determines the *presence* of a *k*-mer, a small SPT compactly specifies the *location* of a *k*-mer. For this work, we consider SPTs where any *k*-mer occurs only once in the set of tiles *𝒰*. The algorithms and ideas introduced in this paper still work with SPTs where a *k*-mer may occur more than once in *𝒰* (some extra book-keeping of a one-to-many *k*-mer-to-tile mapping would be needed, however). For ease of exposition, we ignore tile orientations here. We completely specify the SPT with orientations, allowing tiles to simultaneously represent reverse-complement sequences, in Section S.2.

**Fig. 1.**
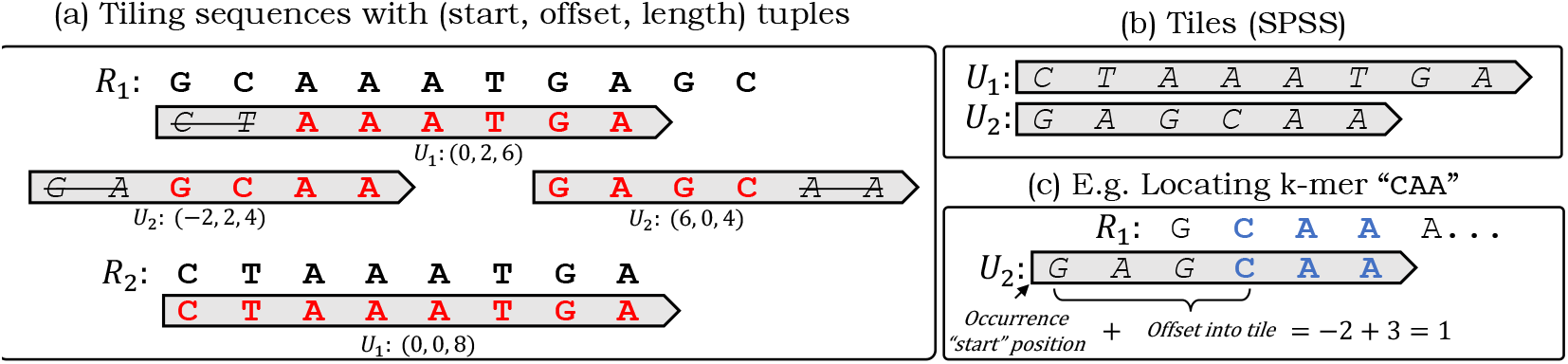
(a) A spectrum preserving tiling (SPT) with *k* = 3, (b) with tiles (an SPSS) that contain all *k*-mers in references. (c) The SPT explicitly encodes where each *k*-mer occurs.

### 3.2 A general and modular index over spectrum preserving tilings

Any SPT is immediately amenable to indexing by an entire *class* of algorithms. This is because an SPT yields a natural decomposition of the MRP query (defined in Section 2) where *k*-mers first map to the tiles and tile-occurrences then map to positions in references. To index a reference collection, a data structure need only compose a query for the positions where *k*-mers occur on tiles in a SPSS with a query for the positions where tiles cover the input references.

Ideally, an index should find a small SPT where *k*-mers are compactly represented in the set of tiles where tiles are “long” and tiling sequences are “short”. Compact tilings exist for almost all practical applications since the amount of *unique* sequence grows much more slowly than the *total* length of reference sequences. Finding a small SPSS where *k*-mers occur only once has been solved efficiently [19,18,20]. However, it remains unclear if a small SPSS induces a small SPT, since an SPT must additionally encode tile-occurrence positions. Currently, tools like pufferfish index reference sequences using an SPT built from the *unitigs* of the compacted de Bruijn graph (cdBG) constructed over the input sequences, which has been found to be sufficiently compact for practical applications. Though the existence of SPSSs smaller than cdBGs suggest that smaller SPTs might be found for indexing, we leave the problem of finding small or even optimal SPTs to future work. Here, we demonstrate how indexing any given SPT is *modular* and possible in general.

Given an SPT, the MRP query can be decomposed into two queries that can each be supported by sparse and efficient data structures. These queries are:

- **The kmer-to-tile query**: Given a *k*-mer *x*, `k2tile`(*x*) returns (*i, p*) — the identity of the tile *U*_*i*_ that contains *x* and the offset (position) into the tile *U*_*i*_ where *x* occurs. That is, `k2tile`(*x*) = (*i, p*) iff *U*_*i*_[*p* : *p* + *k*] = *x*. If *x* is not in *ℛ*, `k2tile`(*x*) returns ∅.
- **The tile-to-occurrence query**: Given the *r*-th occurrence of the tile *U*_*i*_, `tile2occ`(*i, r*) returns the tuple (*n, s, w, l*) that encodes how *U*_*i*_ tiles the reference *R*_*n*_. When `tile2occ`(*i, r*) = (*n, s, w, l*), the *r*-th occurrence of *U*_*i*_ occurs on *R*_*n*_ at position (*s* + *w*), with the sequence *U*_*i*_[*w* : *w*+*l*]. Let the *r*-th occurrence of *U*_*i*_ be *T*_*n,m*_ on *𝒯*, then `tile2occ`(*i, r*) returns (*n, s*_*n,m*_, *w*_*n,m*_, *l*_*n,m*_).

When these two queries are supported, the MRP query can be computed by Algorithm 1. By adding the offset of the queried *k*-mer *x* in a tile *U*_*i*_ to the positions where the tile *U*_*i*_ occurs, Algorithm 1 returns all positions where a *k*-mer occurs. Line 10 checks to ensure that any occurrence of the queried *k*-mer is returned only if the corresponding tile-occurrence of *U*_*i*_ contains that *k*-mer. We note that storing the number of occurrences of a tile and returning `num-occs`(*U*_*i*_) requires negligible computational overhead. In practice, the length of tiling sequences, *𝒯*, are orders of magnitude larger than the number of unique tiles. In this work, we shall use *occ*_*i*_, to denote the number of occurrences of *U*_*i*_ in tiling sequences *𝒯*.

#### Algorithm 1

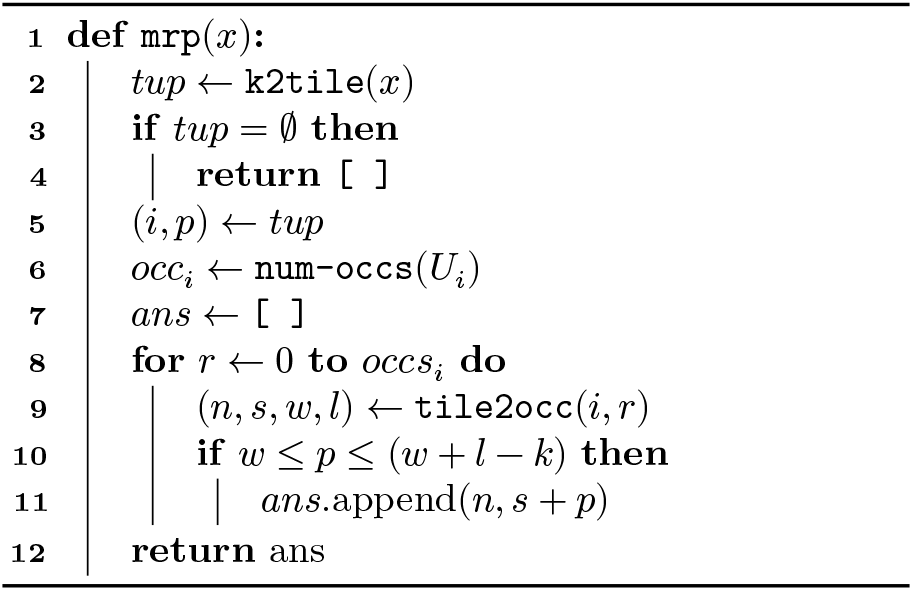

### 3.3 “Drop in” implementations for efficient *k*-mer-to-tile queries

Naturally, prior work for indexing and compressing spectrum preserving string sets (SPSS) can be applied to implement the *k*-mer-to-tile query. When `pufferfish` was first developed, the data structures required to support the *k*-mer-to-tile query dominated the size of moderately sized indexes. Thus, Almodaresi et al. [9] introduced a sampling scheme that samples *k*-mer positions in unitigs. Recently, Pibiri [21,22] introduced `SSHash`, an efficient *k*-mer hashing scheme that exploits minimizer based partitioning and carefully handles highly-skewed distributions of minimizer occurrences. When built over an SPSS, `SSHash` stores the *k*-mers by their order of appearance in the strings (which we term tiles) of an SPSS and thus allows easy computation of a *k*-mer’s offset into a tile. Other methods based on the Burrows-Wheeler transform (BWT) [8], such as the Spectral BWT [23] and BOSS [25], could also be used. However, these methods implicitly sort *k*-mers in lexicographical order and would likely need an extra level of indirection to implement `k2tile`. Unless a compact scheme is devised, this can outweigh the space savings offered by the BWT.

### 3.4 Challenges of the tile-to-occurrence query

The straightforward solution to the tile-to-occurrence query is to store the answers in a table,`utab`, where `utab`[*i*] stores information for all occurrences of the tile *U*_*i*_ and computing `tile2occ`(*i, r*) amounts to a simple lookup into `utab`[*i*][*r*]. This is the approach taken in the pufferfish index and has proven to be effective for moderately sized indexes. This implementation is output optimal and is fast and cache-friendly since all *occ*_*i*_ occurrences of a tile *U*_*i*_ can be accessed contiguously. However, writing down all start positions of tile-occurrences in `utab` is impractical for large indexes.

For larger indexes (e.g. metagenomic references, many human genomes), explicitly storing `utab` becomes more costly than supporting the *k*-mer-to-tile query. This is because, as the number of indexed references grow, the number of distinct *k*-mers grows sub-linearly whereas the number of occurrences grows with the (cumulative) reference length. Problematically, the number of start positions of tile-occurrences grows *at least* linearly. For a reference collection with total sequence length *L*, a naive encoding for `utab` would take *𝒪*(*L* lg *L*) bits, as each position require ⌈lg *L*⌉ bits and there can be at most *L* distinct tiles.

Other algorithms that support “locate” queries suffer from a similar problem. To answer queries in time proportional to the number of occurrences of a query, data structures must explicitly store positions of occurrences and access them in constant time. However, storing *all* positions is impractical for large reference texts or large *k*-mer-sets. To address this, some algorithms employ a scheme to *sample* positions at some small sampling rate *s*, and perform *𝒪*(*s*) work to retrieve not-sampled positions. Since *s* is usually chosen to be a small constant, this extra *𝒪*(*s*) work only imposes a slight overhead.

One may wonder if `utab` — which is an *inverted index* — can be compressed using the techniques developed in the Information Retrieval field [26]. For biological sequences, a large proportion of `utab` consists of very short inverted lists (e.g. unique variants in indexed genomes) that are not well-compressible. In fact, these short lists occur at a rate that is much higher than for inverted indexes designed for natural languages. So, instead applying existing compression techniques, we develop a novel *sampling* scheme for `utab` and the tile-to-occurrence query that exploits the properties of genomic sequences.

## 4 Pufferfish2

Below, we introduce `pufferfish2`, an index built over an SPT consisting of *unitigs*. `Pufferfish2` applies a sampling scheme to sparsify the tile-to-occurrence query of a given `pufferfish` index [9].

### 4.1 Interpreting pufferfish as an index over a unitig-based SPT

Though not introduced this way by Almodaresi et al., `pufferfish` is an index over a *unitig-tiling* of an input reference collection [9]. A *unitig-tiling* is an SPT which satisfies the property that all tiles always occur completely in references where, for every tile-occurrence *T*_*n,m*_ = *U*_*i*_, offset *w*_*n,m*_ = 0 and length *l*_*n,m*_ = |*U*_*i*_|. When this property is satisfied, we term tiles *unitigs*.

An index built over unitig-tilings does not need to store tile-occurrence offsets, *𝒲*, or tile-occurrence lengths *𝔏* since all tiles have the same offset (zero) and occur with maximal length. For indexes constructed over unitig-tilings, we shall use `k2u` to mean `k2tile`, and `u2occ` to be `tile2occ` with one change. That is, `u2occ` omits offsets and lengths of tile occurrences since they are uninformative for unitig-tilings and returns a tuple (*n, s*) instead of (*n, s, w, l*), In prose, we shall refer to these queries as the *k*-mer-to-unitig and unitig-to-occurrence queries.

The MRP query over unitig-tilings can be computed with Algorithm 4 (in Section S.1) where Line 10 is removed from Algorithm 1. We illustrate the MRP query and an example of a unitig-tiling in Fig. 2.

**Fig. 2.**
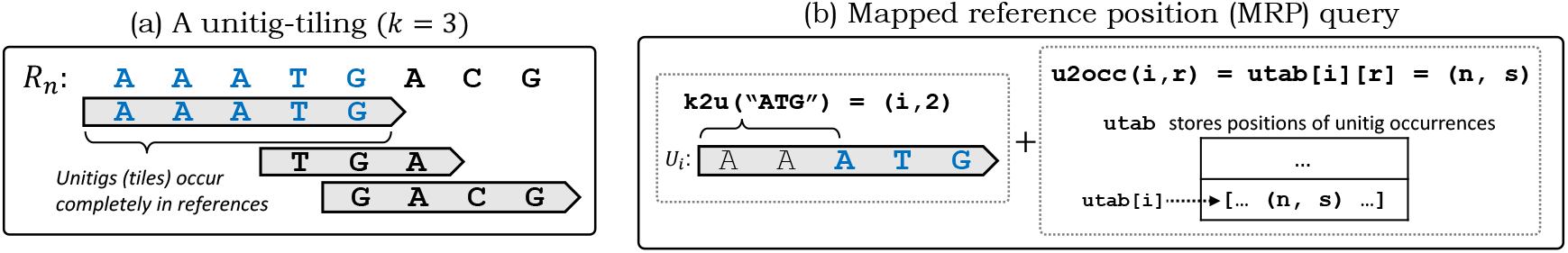
(a) A *unitig-tiling* is an SPT where tiles, *unitigs*, always occur completely in the reference sequences. (b) The MRP query is performed by computing a *k*-mer’s offset into a unitig (`k2u`), then adding the offset to the positions where *unitig-occurrences* appear in indexed reference sequences (`u2occ`). To naively support the unitig-to-occurrence query, positions of all unitig-occurrences are stored in a table, `utab`.

### 4.2 Sampling unitigs and traversing tilings to sparsify the unitig-to-occurrence query

`Pufferfish2` implements a sampling scheme for *unitig-occurrences* on a unitig-tiling. For some small constant *s*, our scheme samples 1/*s* rows in `utab` each corresponding to *all* occurrences of a unique unitig. In doing so, it sparsifies the `u2occ` query and `utab` by only storing positions for a subset of *sampled* unitigs. To compute unitig-to-occurrence queries, it traverses unitig-occurrences on an indexed unitig-tiling.

Notably, `pufferfish2` traverses unitig-tilings that are *implicitly* represented. For unitig-tilings with positions stored in `utab`, there exists no contiguous sequence in memory representing occurrences that is obvious to traverse. However, when viewed as an SPT, *unitig-occurrences* have *ranks* on a tiling and traversals are possible because tiling sequences map uniquely to a sequence of unitig-rank pairs.

Specifically, we define the `pred` query — an atomic traversal step that enables traversals of arbitrary lengths over reference tilings. Given the *r*-th occurrence of the unitig *U*_*i*_, the `pred` query returns the identity and rank of the *preceding* unitig. Let tile *T*_*n,m*_ be the *r*-th occurrence of the unitig *U*_*i*_ on all tiling sequences *𝒯*. Then, `pred`(*i, r*) returns (*j, q*) indicating that *T*_*n,m*−1_, the *preceding* unitig-occurrence, is the *q*-th occurrence of the unitig *U*_*j*_. If there is no preceding occurrence and *m* = 1, `pred`(*i, r*) returns the sentinel value ∅.

When an index supports `pred`, it is able to traverse “backwards” on a unitig-tiling. Successively calling `pred` yields the identities of unitigs that form a tiling sequence. Furthermore, since `pred` returns the identity *j and* the rank *q* of a preceding unitig-occurrence, accessing data associated with each visited occurrence is straightforward in a table like `utab` (i.e., with `utab`[*j*][*q*]).

Given the unitig-set *𝒰*, `pufferfish2` first samples a subset of unitigs *𝒰*_*𝒮*_ ⊆ *𝒰*. For each sampled unitig *U*_*i*_ ∈ *𝒰*_*𝒮*_, it stores information for unitig-occurrences identically to `pufferfish` and records, for *all* occurrences of a sampled unitig *U*_*i*_, a list of reference identity and position tuples in `utab`[*i*].

To recover the position of the *r*-th occurrence a not-sampled unitig *U*_*i*_ and to compute `u2occ`(*i, r*), the index traverses the unitig-tiling and iteratively calls `pred` until an occurrence of a sampled unitig is found — let this be the *q*-th occurrence of *U*_*j*_. During the traversal, `pufferfish2` accumulates number of nucleotides covered by the traversed unitig-occurrences. Since *U*_*j*_ is a sampled unitig, the position of the *q*-th occurrence can be found in `utab`[*j*][*q*]. To return `u2occ`(*i, r*), `pufferfish2` adds the number of nucleotides traversed to the start position stored at `utab`[*j*][*q*], the position of a preceding occurrence of the sampled unitig *U*_*j*_.

This procedure is implemented in Algorithm 2 and visualized in Fig. 3. Traversals must account for (*k* − 1) overlapping nucleotides of unitig-occurrences that tile a reference (Line 5). Storing the length of the unitigs is negligible since the number of unique unitigs is much smaller than the number of occurrences.

**Fig. 3.**
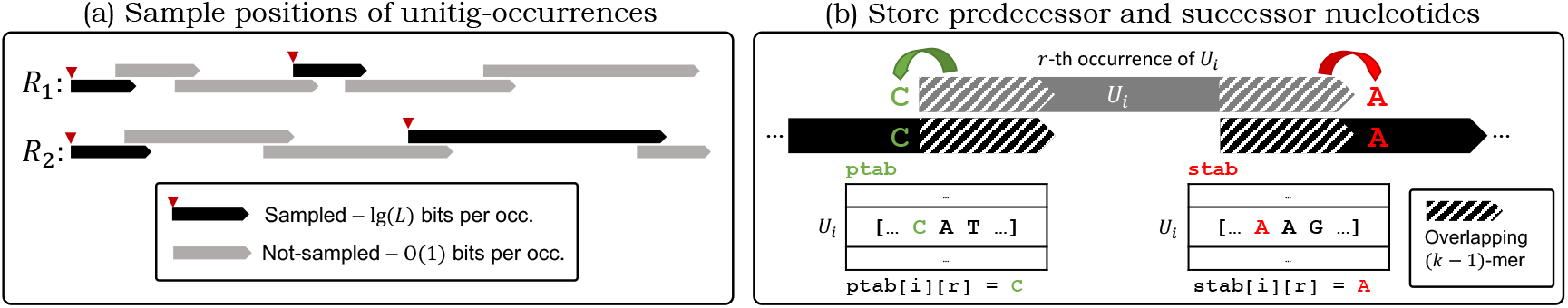
(a) `Pufferfish2` samples unitigs and their occurrences on a unitig-tiling. Only the positions of the occurrences of the *sampled* unitigs (black) are stored in `utab`. Positions of the *not-sampled* unitigs (gray) can be computed relative to the positions of sampled unitigs by traversing backwards on the visualized tiling of references. Sampling the zero-th unitig-occurrence on every reference sequence guarantees that traversals terminate. (b) Predecessor and successor nucleotides are obtained from adjacent unitig occurrences and are stored in the order in which they appear on the references. These nucleotides for the *r*-th occurrence of *U*_*i*_ is stored in `ptab`[*i*][*r*] and `stab`[*i*][*r*], respectively.

#### On the termination of traversals

Any unitig that occurs as the zero-th occurrence (i.e., with rank zero) of a tiling-sequence is always sampled. This way, backwards traversals terminate because every occurrence of a not-sampled unitig occurs after a sampled unitig. This can be seen from Fig. 3. Concretely, if *T*_*n*,1_ = *U*_*i*_ for some tiling-sequence *T*_*n*_, then the unitig *U*_*i*_ must always be sampled.

##### Algorithm 2

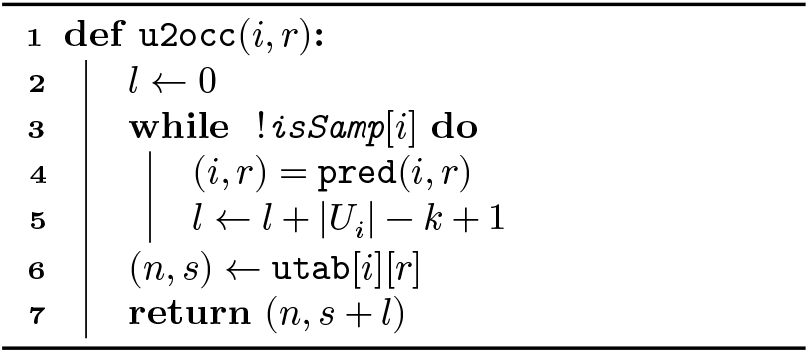

##### Algorithm 3

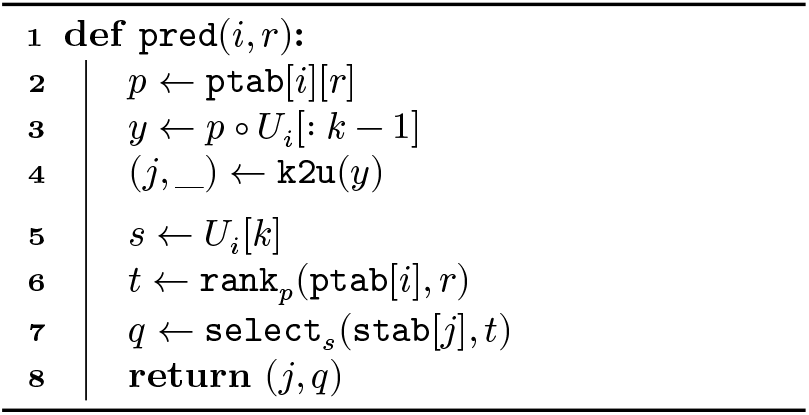

### 4.3 Implementing the pred query with pufferfish2

`Pufferfish2` computes the `pred` query in constant time while requiring only constant space per unitig-occurrence by carefully storing *predecessor* and *successor* nucleotides of unitig-occurrences.

#### Predecessor and successor nucleotides

Given the tiling sequence 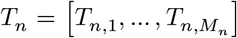, we say that a unitig-occurrence *T*_*n,m*_ is *preceded* by *T*_*n,m*−1_, and that *T*_*n,m*−1_ is *succeeded* by *T*_*n,m*_. Suppose *T*_*n,m*_ = *U*_*i*_, and *T*_*n,m*−1_ = *U*_*j*_, and let the unitigs have lengths *ℓ*_*i*_ and *ℓ*_*j*_, respectively.

We say that, *T*_*n,m*−1_ precedes *T*_*n,m*_ with predecessor nucleotide *p*. The predecessor nucleotide is the nucleotide that precedes the unitig-occurrence *T*_*n,m*_ on the reference sequence *R*_*n*_. Concretely, *p* is the first nucleotide on the last *k*-mer of the preceding unitig, i.e., *p* = *T*_*n,m*−1_[*ℓ*_*j*_ − *k*]. We say that, *T*_*n,m*_ succeeds *T*_*n,m*−1_ with successor nucleotide *s*. Accordingly, the successor nucleotide, *s*, is the last nucleotide on the first *k*-mer of the succeeding unitig, i.e., *s* = *T*_*n,m*_[*k*].

Abstractly, the preceding occurrence *T*_*n,m*−1_ can be “reached” from the succeeding occurrence *T*_*n,m*_ by prepending its predecessor nucleotide to the (*k* − 1)-length prefix of *T*_*n,m*_. Given *T*_*n,m*_ and its predecessor nucleotide *p*, the *k*-mer *Y* that is the last *k*-mer on the preceding occurrence *T*_*n,m*−1_ can be obtained with *Y* = *p* ∘ *T*_*n,m*_[: *k* − 1]. Given an occurrence *T*_*n,m*_, let the functions `pred-nuc` (*T*_*n,m*_) and `succ-nuc` (*T*_*n,m*_) yield the predecessor nucleotide and the successor nucleotide of *T*_*n,m*_, respectively. If *T*_*n,m*_ is the first or last unitig-occurrence pair on *T*_*n*_, then `succ-nuc` (*T*_*n,m*_) and `pred-nuc` (*T*_*n,m*_) return the “null” character, ‘$’.

These notationally dense definitions can be more easily understood with a figure. Figure 3 shows how predecessor and successor nucleotides of a given unitig-occurrence on a tiling are obtained.

#### Concrete representation

`Pufferfish2` first samples a set of unitigs *𝒰*_*𝒮*_ ⊆ *𝒰* from *𝒰* and stores a bit vector, `isSamp`, to record if a unitig *U*_*i*_ is sampled where `isSamp`[*i*] = 1 iff *U*_*i*_ ∈ *𝒰*_*𝒮*_. `Pufferfish2` stores in `utab` the reference identity and position pairs for occurrences of *sampled* unitigs only.

After sampling unique unitigs, `pufferfish2` stores a *predecessor nucleotide table*, `ptab`, and a *successor nucleotide table*, `stab`. For each not-sampled unitig *U*_*i*_ *only*, `ptab`[*i*] stores a list of predecessor nucleotides for each occurrence of *U*_*i*_ in the unitig-tiling. For *all* unitigs *U*_*i*_, stab[*i*] stores a list of successor nucleotides for each occurrence of *U*_*i*_. Concretely, when the unitig-occurrence *T*_*n,m*_ is the *r*-th occurrence of *U*_*i*_,

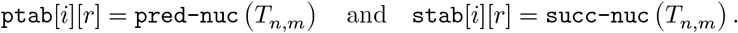

As discussed in Section 4.2, unitigs that occur as the zero-th element on a tiling is always sampled so that every occurrence of a not-sampled unitig has a predecessor. If *T*_*n,m*_ has no successor and is the last unitig-occurrence on a tiling sequence, `stab`[*i*][*j*] contains the sentinel symbol ‘$’. Figure 3 illustrates how predecessor and successor nucleotides are stored.

#### Computing the pred query

Given the *k*-mer-to-unitig query, `pufferfish2` supports the `pred` query for any unitig *U*_*i*_ that is not-sampled. When the *r*-th occurrence of *U*_*i*_ succeeds the *q*-th occurrence of *U*_*j*_, it computes `pred`(*i, r*) = (*j, q*) with Algorithm 3. To compute `pred`, it constructs a *k*-mer to find *U*_*j*_, and then computes one rank and one select query over the stored lists of nucleotides to find the correct occurrence.

`Pufferfish2` first computes *j*, the identity of the preceding unitig. The last *k*-mer on the preceding unitig must be the first (*k* − 1)-mer of *U*_*i*_ *prepended* with predecessor nucleotide of the *r*-th occurrence of *U*_*i*_. Given ptab[*i*][*r*] = *p*, it constructs the *k*-mer, *Y* = *p* ∘ *U*_*i*_[: *k* − 1], that must be the last *k*-mer on *U*_*j*_. So on Line 4, it computes `k2u`(*Y*) to obtain the identity of the preceding unitig *U*_*j*_.

It then computes the unitig-rank, *q*, of the preceding unitig-occurrence of *U*_*j*_. Each time *U*_*i*_ is preceded by the nucleotide *p*, it must be preceded by the *same* unitig *U*_*j*_ since any *k*-mer occurs in only one unitig. Accordingly, each occurrence *U*_*j*_ that is succeeded by *U*_*i*_ must always be succeeded by the *same* nucleotide *s* equal to the *k*-th nucleotide of *U*_*i*_, *U*_*i*_[*k*]. For the preceding occurrence of *U*_*j*_ that the algorithm seeks to find, the nucleotide *s* is stored at some unknown index *q* in `stab`[*j*] — the list of successor nucleotides of *U*_*j*_.

Whenever an occurrence of *U*_*i*_ succeeds an occurrence of *U*_*j*_, so do the corresponding pair predecessor and successor nucleotides stored in `ptab`[*i*] and `stab`[*j*]. Since `ptab`[*i*] and `stab`[*j*] store predecessor and successor nucleotides in the order in which unitig-occurrences appear in the tiling sequences, the following *ranks* of stored *nucleotides* must be equal: (1) the rank of the nucleotide *p* = `ptab`[*i*][*r*] at index *r* in the list of predecessor nucleotides, `ptab`[*i*], of the succeeding unitig *U*_*i*_, and (2) the rank of the nucleotide *s* = *U*_*i*_[*k*] at index *q* in the list of successor nucleotides, `stab`[*j*], of the preceding unitig *U*_*j*_. We illustrate this correspondence between ranks in Fig. 4. So to find *q*, the rank of the preceding unitig-occurrence, `pufferfish2` computes the rank of the predecessor nucleotide, *t* = rank_*p*_(ptab[*i*], *r*). Then, computing `select`_*s*_(`stab`[*i*], *t*), the index where the *t*-th rank successor nucleotide of *U*_*j*_ occurs must yield *q*.

**Fig. 4.**
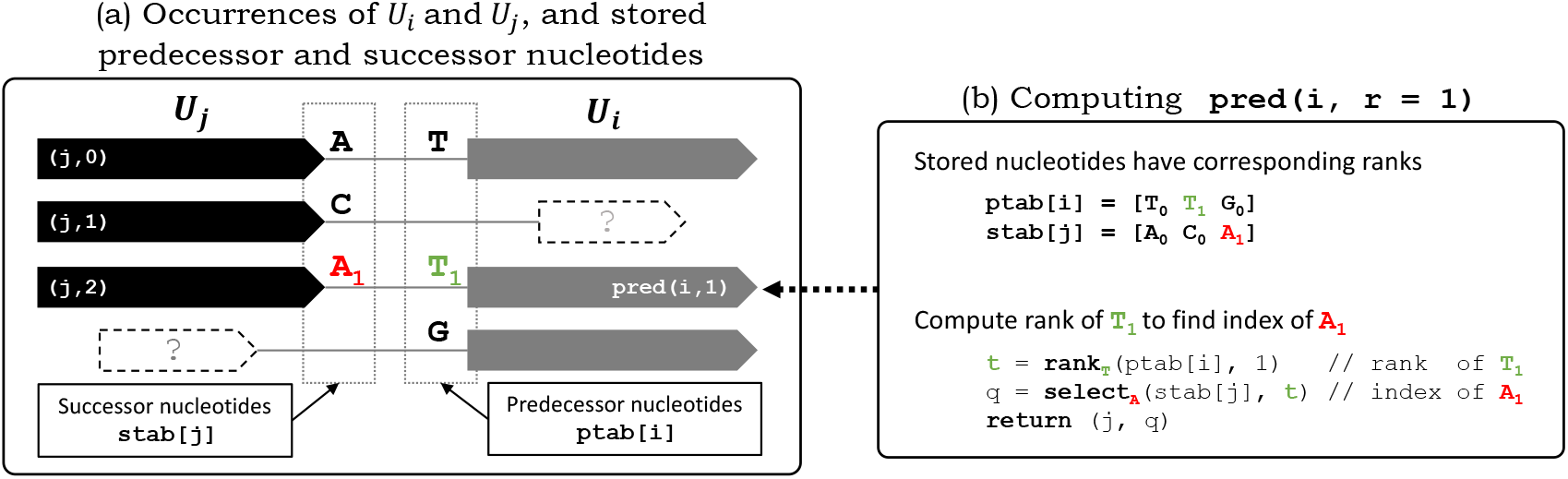
Visualizing the `pred` query that finds the occurrence of *U*_*j*_ that precedes the queried occurrence of *U*_*i*_ with rank 1. (a) All occurrences of *U*_*i*_ and *U*_*j*_ are visualized (in sorted order) with their preceding and succeeding unitig occurrences, respectively. The figure shows stored successor nucleotides for *U*_*j*_, and predecessor nucleotides for *U*_*i*_. Whenever an occurrence of *U*_*j*_ precedes an occurrence of *U*_*i*_, a corresponding pair of nucleotides `“A”` and `“T”` occur and are stored in `stab`[*j*] and `ptab`[*i*] respectively. (b) Their *ranks* (annotated with subscripts) of the corresponding predecessor-successor nucleotide pair *match* in `ptab`[*i*] and `stab`[*j*], but the *indices* do not. A rank query for predecessor nucleotide `“T”` at index *r* = 1 yields the matching rank of the successor nucleotide `“A”`. A select query for the nucleotide `“A”` with rank 1 yields the *index* and occurrence of the predecessor *U*_*j*_.

#### Time and space analysis

`Pufferfish2` computes the `pred` query in constant time. The *k*-mer for the query k2u is assembled in constant time, and the k2u query itself is answered in constant time, as already done in the `pufferfish` index [9].

For not-sampled unitigs, `pufferfish2` does not store positions of unitig-occurrences in `utab`. Instead, it stores nucleotides in tables `stab` and `ptab`. These tables are implemented by *wavelet matrices* that support rank, select, and access operations in *𝒪*(lg *σ*) time on sequences with alphabet size *σ* while requiring only lg *σ* + *o*(lg *σ*) bits per element [24].

As explained in Section 3.1, we have avoided the treatment of *orientations* of nucleotide sequences for brevity. In actuality, unitigs may occur in a *forward* or a *backwards* orientation (i.e., with a reverse complement sequence). When considering orientations, `pufferfish2` implements the `pred` query by storing and querying over lists of *nucleotide-orientation* pairs. In this case, `ptab` and `stab` instead store predecessor-orientation and successor-orientation pairs. Accordingly, wavelet matrices are then built over alphabets of size 8 and 9 respectively — deriving from eight nucleotide-orientation pairs and one sentinel value for unitig-occurrences that have no predecessor. Thus, `ptab` and `stab` in total require ≈ 7 bits per unitig-occurrence (since 7 = ⌈lg 8⌉ +⌈lg 9⌉). We describe how the `pred` query is implemented with orientations in Section S.3.

#### Construction

The current implementation of `pufferfish2` sparsifies the unitig-to-occurrence query and compresses the table of unitig occurrences, `utab`, of an existing `pufferfish` index, and inherits its *k*-mer-to-unitig mapping. In practice, sampling and building a `pufferfish2` index always takes less time than the initial `pufferfish` index construction. In brief, building `pufferfish2` amounts to a linear scan over an SPT. We describe how `pufferfish2` in constructed in more detail in Section S.4.

### 4.4 A random sampling scheme to guarantee short backwards traversals

Even with a constant-time `pred` query, computing the unitig-to-occurrence query is fast only if the length of backwards traversals — the number of times `pred` is called — is small. So for some small constant *s*, a sampling scheme should sample 1/*s* of *unique* unitigs, store positions of only 1/*s* of unitig-*occurrences* in `utab`, and result in traversal lengths usually of length *s*.

At first, one may think that a greedy sampling scheme that traverses tiling sequences to sample unitigs could be used to bound traversal lengths to some given maximum length, *s*. However, when tiling sequences become much longer than the number of unique unitigs, such a greedy scheme samples almost *all* unitigs and only somewhat effective in limited scenarios (see Section S.5). Thus, we introduce the *random* sampling scheme that samples 1/*s* of unitigs uniformly at random from *𝒰*. This scheme guarantees that traversals using the `pred` query terminate in *s* steps *in expectation* if each unitig-occurrence *T*_*n,m*_ is independent and identically distributed and drawn from an arbitrary distribution. Then, backwards traversals until the occurrence of a sampled unitig is a series of Bernoulli trials with probability 1/*s*, and traversal lengths follow a geometric distribution with mean *s*. Although this property relies on a simplifying assumption, the random sampling scheme works well in practice.

### 4.5 Closing the gap between a constant time pred query and contiguous array access

Even though the `pred` query is constant time and traversals are short, it is difficult to implement `pred` queries in with speed comparable to *contiguous array accesses* that are used to compute the `u2occ` for when `utab` is “dense” — i.e., uncompressed and not sampled. In fact, any compression scheme for `utab` would have difficulty contending with constant time contiguous array access regardless of their asymptotics since dense implementations are output optimal, very cache friendly, and simply store the answers to queries in an array. To close the gap between theory and practice, `pufferfish2` exploits several optimizations.

In practice, a small proportion of unique unitigs are “popular” and occur extremely frequently. Fortunately, the total number of occurrences of popular unitigs is small relative to other unitigs. To avoid an excessively large number of traversals from a not-sampled unitig, `pufferfish2` modifies the sampling scheme to always sample popular unitigs that occur more than a preset number, *α*, times. Better yet, we re-parameterize this optimization and set *α* so that the total number of occurrences of popular unitigs sum to a given proportion 0 < *t* ≤ 1 of the total occurrences of all the unitigs. For example, setting *t* = 0.25 restricts `pufferfish2` to sample from 75% of the total size of `utab` consisting of unitigs that occur most infrequently.

Also, the MRP and `pred` query are especially amenable to caching. Notably, `pufferfish2` caches and memoizes redundant `k2u` queries in successive `pred` queries. Also, it caches “streaming” queries to exploit the fact that successive queried *k*-mers (e.g., from the same sequenced read) likely land on the same unitig. We describe in more detail these and other important optimizations in Section S.6.

## 5 Experiments

We assessed the space-usage of the indexes constructed by `pufferfish2` from several different wholegenome sequence collections, as well as its query performance with different sampling schemes. Reported experiments were performed on a server with an Intel Xeon CPU (E5-2699 v4) with 44 cores and clocked at 2.20 GHz, 512 GB of memory, and a 3.6 TB Toshiba MG03ACA4 HDD.

### Datasets

We evaluated the performances on a number of datasets with varying attributes: (1) Bacterial collection: a random set of 4000 bacterial genomes from the NCBI microbial database; (2) Human collection: 7 assembled human genome sequences from [27]; and (3) Metagenomic collection: 30,691 representative sequences from the most prevalent human gut prokaryotic genomes from [28].

### Results

To emulate a difficult query workload, we queried the indexes with 10 million random *true positive k*-mers sampled uniformly from the indexed references. Our results from Table 1 show that sampling *popular* unitigs is critical to achieve reasonable trade-offs between space and speed. When indexing seven human genomes, the difference in space between always sampling using *t* = 0.05 and *t* = 0.25, is only 2.1GB (12.5% of the uncompressed `utab`). However, explicitly recording 2.1GB of positions of occurrences of popular unitigs, *substantially* reduces the comparative slowdown from 43.8× to 7.9×. This is because setting *t* = 0.25 instead of *t* = 0.05 greatly reduces the maximum number of occurrences of a *not-sampled* unitig — from ≈87,000 to ≈9,000 times, respectively. Here, setting *t* = 0.25 means that random *k*-mer queries that land in not-sampled unitigs perform many fewer traversals over reference tilings.

**Table 1.** Size and speed of `pufferfish2` indexes querying 10 million random *k*-mers and 100,000 reads. Uncompressed, baseline implementations of the unitig-to-occurrence query (`pufferfish` indexes with the *sparse* `k2u` implementation [9]) are labeled with “None” sampling strategy. Relative sizes of compressed representations and relative slowdowns to the baseline are indicated in parentheses.

| Dataset | Sampling strategy | u2occ size (GB) | 10M $k$-mers (secs) | 100K reads (secs) |
| --- | --- | --- | --- | --- |
| 7 Humans | None | 16.8 | 86.1 | 139.4 |
| | Random ($s = 3, t = .05$) | 7.8 (0.46) | 4159.1 (43.8×) | 8092.8 (58.04×) |
| | Random ($s = 3, t = .25$) | 9.9 (0.59) | 681.1 (7.9×) | 1466.2 (10.52×) |
| 4000 Bacteria | None | 7.7 | 35.5 | 12.6 |
| | Random ($s = 3, t = .05$) | 3.7 (0.48) | 420.4 (11.9×) | 15.6 (1.24×) |
| | Random ($s = 3, t = .25$) | 4.7 (0.61) | 323.8 (9.1×) | 15.5 (1.23×) |
| 30K Human gut | None | 86.3 | 80.6 | 178.7 |
| | Random ($s = 3, t = .05$) | 45.6 (0.53) | 439.4 (5.5×) | 570.2 (3.19×) |
| | Random ($s = 3, t = .25$) | 54.4 (0.63) | 365.2 (4.5×) | 576.9 (3.23×) |
| | Random ($s = 6, t = .05$) | 34.6 (0.40) | 1037.5 (12.9×) | 644.8 (3.61×) |
| | Random ($s = 6, t = .25$) | 45.6 (0.53) | 614.0 (7.6×) | 646.1 (3.56×) |

On metagenomic datasets, indexes are compressed to a similar degree but differences in query speed at different parameter settings are small. `Pufferfish2` is especially effective for a *large* collection of bacterial genomes. With the fastest parameter setting, it incurs only a 4.5× slowdown for random queries while reducing the size of `utab` for the collection of 30,000 bacterial genomes by 37% (from 86.3GB to 54.4GB).

Apart from random lookup queries, we also queried the indexes with *k*-mers deriving from sequenced readsets [29,30]. We measured the time to query and recover the positions of all *k*-mers on 100,000 reads. This experiment demonstrates how the slowdown incurred from sampling can (in most cases) be further reduced when queries are positionally coherent or miss. Successive *k*-mer queries from the same read often land on the same unitig and can thus be cached (see Section 4.5). *True negative k*-mers that do not occur in the indexed reference collection neither require traversals nor incur any slowdowns.

To simulate a metagenomic analysis, we queried reads from a human stool sample against 4,000 bacterial genomes. This is an example of a low hit-rate analysis where 18% of queried *k*-mers map to indexed references. In this scenario, `pufferfish2` reduces the size of `utab` by *half* but incurs only a 1.2× slowdown. We also queried reads from the same human stool sample against the collection of 30,000 bacterial genomes representative of the human gut. Here, 88% of *k*-mers are found in the indexed references. At the sparsest setting, `pufferfish2` indexes incur only a 3.6× slowdown while reducing the size of `utab` by 60%.

We observe that `pufferfish2’s` sampling scheme is less effective when indexing a collection of seven human genomes. When sampled with *s* = 3 and *t* = 0.25, `pufferfish2` incurs a 10.5× slowdown when querying reads from a DNA-seq experiment in which 92% of queried *k*-mers occur in reference sequences. Interestingly, the slowdown when querying reads is larger than the slowdown when querying random *k*-mers. This is likely due to biases from sequencing that cause *k*-mers and reads to map to non-uniformly indexed references. Nonetheless, this result motivates future work that could design sampling schemes optimized for specific distributions of query patterns.

We expect to see less-pronounced slowdowns in practice than those reported in Table 1. This is because tools downstream of an index like `pufferfish2` almost always perform operations *much* slower after straightforward exact lookups for *k*-mers. For example, aligners have to perform alignment accounting for mismatches and edits. Also, our experiments pre-process random *k*-mer sets and read-sets so that no benchmark is I/O bound. Critically, the compromises in speed that `pufferfish2` makes are especially palatable because it trades-off speed in the *fastest* operations in analyses — *exact k*-mer queries — while substantially reducing the space required for the *most space intensive* operation.

### Using SSHash for even smaller indexes

For convenience, we have implemented our SPT compression scheme within an index that uses the *specific* sparse `pufferfish` implementation for the *k*-mer-to-tile (*k*-mer-to-unitig) mapping [9]. However, the SPT enables the construction of modular indexes that use *various* data structures for the *k*-mer-to-tile mapping and the tile-to-reference mapping, provided only a minimalistic API between them. A recent representation of the *k*-mer-to-tile mapping that supports all the necessary functionality is `SSHash` [22]. Compared to the k2u component of `pufferfish, SSHash` is almost always substantially smaller. Further, it usually provides faster query speed compared to the *sparse* `pufferfish` implementation of the *k*-mer-to-tile query, especially when streaming queries are being performed.

In Table 2, we calculate the size of indexes if `SSHash` is used for the *k*-mer-to-tile mapping — rather than the *sparse* `pufferfish` implementation. These sizes then represent overall index sizes that would be obtained by pairing a state-of-the-art representation of the *k*-mer-to-tile mapping with a state-of-the-art representation of the tile-to-reference mapping (that we have presented in this work). Practically, the only impediment to constructing a fully-functional index from these components is that they are implemented in different languages (`C++` for `SSHash` and Rust for `pufferfish2`) — we are currently addressing this issue.

**Table 2.** Sizes in GB of possible, new indexes — with `k2u` implemented by `SSHash` and `u2occ` by `pufferfish2` — compared to the size of original `pufferfish` indexes. Selected sampling parameters for datasets (top-to-bottom) are (*s* = 3, *t* = 0.25), (*s* = 3, *t* = 0.05), and (*s* = 6, *t* = 0.05), respectively.

| Dataset | u2occ w/ **pufferfish2** | k2u w/ **SSHash** | New index | Original **pufferfish** index |
| --- | --- | --- | --- | --- |
| 7 Human | 9.9 | 3.2 | **13.1** | 28.0 |
| 4000 Bacteria | 3.7 | 7.3 | **11.0** | 26.1 |
| 30K Human gut | 34.6 | 22.0 | **55.6** | 131.7 |

Importantly, these results demonstrate that, when `SSHash` is used, the representation of the tile-to-occurrence query becomes a bottleneck in terms of space, occupying an increasingly larger fraction of the overall index. Table 2 shows that, in theory, if one fully exploits the modularity of SPTs, new indexes that combine `SSHash` with `pufferfish2` would be *half* the space of the original pufferfish index. As of writing, with respect to an index over 30,000 bacterial genomes, the estimated difference in *monetary* cost of an AWS EC2 instance that can fit a new 55.6GB index versus a 131GB `pufferfish` index in memory is 300USD per month (see Section S.7).

### Comparing to MONI and the r-index

We compared `pufferfish2` to MONI, a tool that builds an r-index to locate maximal exact matches in highly repetitive reference collections [6]. In brief, `pufferfish2` is faster and requires less space than MONI for our benchmarked bacterial dataset. Our tool does so with some trade-offs. `Pufferfish2` supports rapid locate queries for *k*-mers of a *fixed* length, while r-index based approaches supports locate queries for patterns of any *arbitrary* length and can be used to find MEMs. Notably, it has been shown that both *k*-mer and MEM queries can be used for highly effective read-mapping and alignment [1,6].

For reference, we built MONI on our collection of 4,000 bacterial genomes. Here, MONI required 51.0G of disk space to store which is 29% larger than the `pufferfish` index (39.5GB) with its *dense* k2u implementation — its *least* space-efficient configuration. The most space efficient configuration of the `pufferfish2` index (with *s*=3, *t*=.25) is 42% the size of MONI when built on from the same data and requires 21.7GB of space. Compared to a theoretically possible index specified in Table 2 that would only require 11.0GB, MONI would need 4.6× more space.

We also performed a best-effort comparison of query speed between `pufferfish2` and MONI. Unfortunately, it is not possible to directly measure the speed of exact locate queries for MONI because it does not expose an interface for such queries. Instead, we queried MONI to find MEMs on true-positive *k*-mers treating each *k*-mer as unique read (encoded in FASTQ format as MONI requires). We argue that this is a reasonable proxy to exact locate queries because, for each true-positive *k*-mer deriving from an indexed reference sequence, the entire *k*-mer itself is the maximal exact match. For MONI, just like in benchmarks for in Table 1, we report the time taken for computing queries only and ignore time required for I/O operations (i.e. loading the index and quries, and writing results to disk).

We found that `pufferfish2` is faster than MONI when querying *k*-mers against our collection of 4,000 bacterial genomes. MONI required 1,481.7 seconds to query the same set of 10 million random true-positive k-mers queried in Table 1. When compared to the slowest built most space efficient configuration of `pufferfish2` benchmarked in Table 1, `pufferfish2` is 3.5× faster.

## 6 Discussion and future work

In this work, we introduce the *spectrum preserving tiling* (SPT), which describes how a spectrum preserving string set (SPSS) tiles and “spells” an input collection of reference sequences. While considerable research effort has been dedicated to constructing space and time-efficient indexes for SPSS, little work has been done to develop efficient representations of the tilings themselves, despite the fact that these tilings tend to grow more quickly than the SPSS and quickly become the size bottleneck when these components are combined into reference indexes. We describe and implement a sparsification scheme in which the space required for representing an SPT can be greatly reduced in exchange for an expected constant-factor increase in the query time. We also describe several important heuristics that are used to substantially lessen this constant-factor in practice. Having demonstrated that modular reference indexes can be constructed by composing a *k*-mer-to-tile mapping with a tile-to-occurrence mapping, we have thus opened the door to exploring an increasingly diverse collection of related reference indexing data structures.

Despite the encouraging progress that has been made here, we believe that there is much left to be explored regarding the representation of SPTs, and that many interesting questions remain open. Some of these questions are: (1) How would an algorithm sample individual unitig-occurrences instead of all occurrences of a unitig to *explicitly* bound the lengths of backwards traversals? (2) Does a smaller SPSS imply a small SPT and could one compute an optimally small SPT? (3) Given some distributional assumptions for queries, can an algorithm sample SPTs to minimize the expected query time? (4) In practice, how can an implemented tool combine our sampling scheme with existing compression algorithms for the highly skewed tile-to-occurrence query? (5) Can a *lossy* index over an SPT be constructed and applied effectively in practical use cases?

With excitement, we discuss in more detail these possibilities for future work in more detail in Section S.8.

## Supplementary materials for

### S.1 The mapped position query (MRP) for unitig-tilings

#### Algorithm 4

The MRP query for unitig-tilings

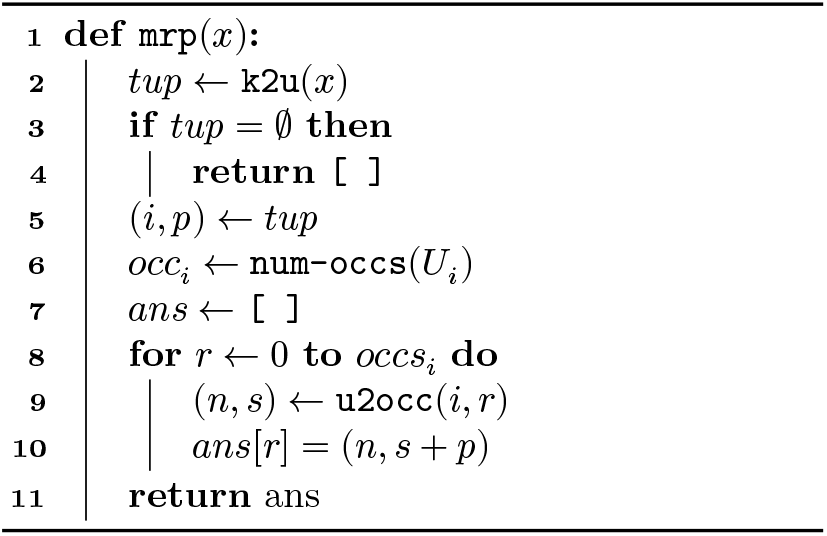

### S.2 Spectrum preserving tilings with *orientations*

We extend the definition of spectrum preserving tilings (without orientations) given in Section 3.1, to formally define spectrum preserving tilings (SPT) *with orientations*. An SPT with orientation allows tiles (members of a spectrum preserving string set) to occur in either a *forward* orientation as stored in memory as a nucleotide sequence, or a *backwards* orientation as the *reverse complement* of the stored sequence.

With respect to representing reference genomic sequences, using SPTs with orientations is particularly useful because it avoids redundantly encoding and storing occurrences of a *k*-mer *and* the reverse complement of said *k*-mer. Furthermore, since most sequencing technologies are agnostic to strands of DNA sequences, considering orientations enables the simultaneous and canonical representation of both corresponding strands of an indexed genomic sequence.

Also, as in [9], we consider only *odd k*-mer sizes so that no *k*-mer is its own reverse complement.

#### Tiling sequences of tile *and orientation* pairs

Given a fixed *k*-mer size, *k*, a tiling sequence *T*_*n*_ in *𝒯* is instead sequences of *tile-orientation* pairs where each occurrence is defined to be *T*_*n,m*_ = (*U*_*i*_, *o*), for some unitig *U*_*i*_ ∈ *𝒰* and an orientation *o* ∈ {0, 1}. Here, *o* = 1 indicates that the unitig *U*_*i*_ occurs in a forward orientation and *o* = 0 indicates that it occurs in the *backwards* orientation with reverse complement sequence *Ū*_*i*_. Notationally, *Ū*_*i*_ is the string that is the reverse complement of *U*_*i*_ where *U*_*i*_ is reversed and each nucleotide is replaced with its complement.

Let us define the `spell`(*U*_*i*_, *o*) function for a unitig orientation pair to return the forward sequence *U*_*i*_ if *o* = 1 and the backwards, reverse complement sequence *Ū*_*i*_ otherwise. Abusing some notation, when *T*_*n,m*_ = (*U*_*i*_, *o*), let `spell`(*T*_*n,m*_) = `spell`(*U*_*i*_, *o*).

Then formally, a spectrum preserving tiling with orientations for a reference collection *ℛ* = {*R*_1_, …, *R*_*N*_ } tiles each reference sequence *R*_*n*_ with sequences *spelled by* occurrences of tile-orienation pairs. Specifically, each *R*_*n*_ can be reconstructed by gluing together the sequences that tile-orientation pairs *spell*. For each *R*_*n*_ that is tiled by *M*_*n*_ tile occurrences, the SPT satisfies the property that:

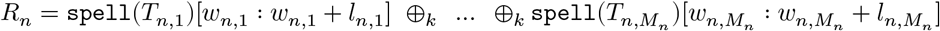

#### S.2.1 Returning queries with orientations

Accordingly, when indexing an SPT with orientations the mapped reference position query, *k*-mer-to-tile query, and tile-to-occurrence query also return orientations. Here, we extend and reintroduce the queries defined in Section 3.

1. **The mapped reference position (MRP) query** Given any *k*-mer *x*, the MRP query enumerates the positions and orientations of all occurrences of *x* in *ℛ*. Precisely, each returned occurrence is a tuple (*n, p, o*), that specifies that *k*-mer *x* occurs in reference *n* at position *p* with orientation *o*. That is, if *o* = 1, then *x* occurs in the forward orientation as *R*_*n*_[*p* : *p*+*k*] = *x*. Otherwise, the reverse complement occurs as 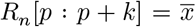. If a *k*-mer does not occur in some *R*_*n*_ ∈ *ℛ*, the query returns an empty list.
2. **The kmer-to-tile query**: Given a *k*-mer *x*, `k2tile`(*x*) returns (*i, p, o*) — the identity of the tile *U*_*i*_ that contains *x*, the offset (position) into the tile *U*_*i*_ where *x* occurs, and the *orientation* of how *x* occurs. That is, `k2tile`(*x*) = (*i, p*, 1) if *U*_*i*_[*p* : *p* + *k*] = *x*, and `k2tile`(*x*) = (*i, p*, 0) if 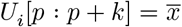 where *x* occurs in the backwards orientation as the reverse complement. If *x* is not in *ℛ*, `k2tile, k2tile`(*x*) returns ∅.
3. **The tile-to-occurrence query**: Given the *r*-th occurrence of the tile *U*_*i*_, tile2occ(*i, r*) returns the tuple (*n, o, s, w, l*) that encodes how *and in what orientation U*_*i*_ tiles the reference *R*_*n*_. Let the *r*-th occurrence of *U*_*i*_ be a tile-occurrence *T*_*n,m*_ on *𝒯* where *T*_*n,m*_ = *U*_*i*_, *o* for some orientation *o*. Then `tile2occ`(*i, r*) returns (*n, o, s*_*n,m*_, *w*_*n,m*_, *l*_*n,m*_). When `tile2occ`(*i, r*) = (*n, o, s, w, l*) and *o* = 1, the *r*-th occurrence of *U*_*i*_ occurs on *R*_*n*_ at position (*s* + *w*), with the sequence *U*_*i*_[*w* : *w* + *l*]. When `tile2occ`(*i, r*) = (*n, o, s, w, l*) and *o* = 0, the *r*-th occurrence of *U*_*i*_ occurs on *R*_*n*_ at position (*s* + *w*), with the sequence *Ū*_*i*_[*w* : *w* + *l*].

With some arithmetic bookkeeping considering orientations and lengths, the MRP query with orientations can again be decomposed into the two corresponding *k*-mer-to-tile and tile-to-occurrence queries that also return orientations. Although not introduced with respect to an SPT, the pufferfish index developed by Almodaresi et al. [9] is implemented exactly this way as an index over an SPT *with orientations* of unitigs.

### S.3 Pufferfish2: the pred query with orientations

`Pufferfish2’s` sampling scheme and the `pred` query can be applied when considering orientations — our implemented tool does exactly this. Below, we extend Section 4 to fully specify the introduced sampling scheme and the `pred` query when orientations are considered.

#### S.3.1 Predecessor and successor nucleotides

When SPT references, predecessor and successor nucleotides are defined and obtained with respect to sequences on the *references*. Specifically, the predecessor nucleotide is the first nucleotide of the last *k*-mer on of the preceding unitig-occurrence *as spelled with the corresponding orientation* of the occurrence. The successor nucleotide is defined in the same manner.

Suppose *T*_*n,m*_ = (*U*_*i*_, *o*), and *T*_*n,m*−1_ = (*U*_*j*_, *ω*), and let the unitigs have lengths *ℓ*_*i*_ and *ℓ*_*j*_, respectively. We say that, *T*_*n,m*−1_ precedes *T*_*n,m*_ with predecessor nucleotide *p and orientation o*. Concretely, *p* is the first nucleotide on the last *k*-mer of the preceding unitig, with *p* = `spell`(*T*_*n,m*−1_)[*ℓ*_*j*_− *k*]. We say that, *T*_*n,m*_ succeeds *T*_*n,m*−1_ with successor nucleotide *s and orientation ω*. Accordingly, the successor nucleotide, *s*, is the last nucleotide on the first *k*-mer of the succeeding unitig, with *s* = `spell`(*T*_*n,m*_)[*k*].

#### S.3.2 Storing nucleotide-orientation pairs in ptab and stab

Instead of storing only nucleotides, `pufferfish2` stores nucleotide-orientation pairs in implementation. That is, for each occurrence *T*_*n,m*_ = (*U*_*i*_, *o*) that is the *r*-th occurrence of a not-sampled unitig *U*_*i*_,

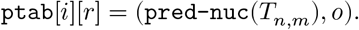

And for each occurrence *T*_*n,m*_ = (*U*_*i*_, *o*) that is the *r*-th occurrence of *any* unitig *U*_*i*_,

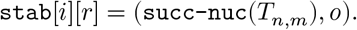

In summary, `ptab` and `stab` store for each corresponding unitig-occurrence, the nucleotides that succeed and precede it as they occur on a tiled reference, *and* the orientation of said occurrence.

#### S.3.3 Computing the pred query by matching ranks of predecessor-orientation and successor-orientation pairs

When orientations are considered, computing the `pred` query requires matching ranks of predecessor-orientation and successor-orientation pairs. Critically, any time a pair of unitigs occur as a successor-predecessor pair in *fixed* orientations, the corresponding pair of predecessor and successor nucleotides are *consistent* and also *fixed*. Furthermore, if an occurrence of the *U*_*j*_ in orientation *ω* precedes a unitig *U*_*i*_ with orientation *o, any other* occurrence of *U*_*j*_ that precedes *U*_*i*_ with orientation *o* must also occur with orientation *ω*. We state and prove this property with Theorem 1 and illustrate examples of both possible and impossible unitig-occurrences with Fig. S1.

Theorem 1 guarantees that whenever *U*_*i*_ occurs with orientation *o* with predecessor nucleotide *p* preceded by *U*_*j*_, *U*_*j*_ must occur with fixed orientation *ω* with a fixed successor nucleotide *s*. We illustrate this correspondence in Fig. S1. Algorithm 5 implements pred with orientations considered. To find the identity, *j*, of the preceding unitig occurrence, Algorithm 5 must construct the last

*k*-mer of the corresponding occurrence *U*_*j*_ as it appears on the reference. Specifically, in Line 3 it spells *U*_*i*_ before extracting the overlapping (*k* − 1)-mer. Here, k2u returns orientation, *ω*, of the queried *k*-mer on *U*_*j*_, which must also be the orientation of the preceding unitig occurrence on the reference. Furthermore, the successor nucleotide, *s*, for the preceding occurrence of *U*_*j*_ must be the *k*-th nucleotide on *U*_*i*_ spelled with orientation *o* (Line 5).

Now, Algorithm 5 has all it needs to compute *q*, the unitig-rank of the preceding occurrence of *U*_*j*_. Computing the rank of (*p, o*) in ptab[*i*] yields the rank of the corresponding successor-orientation pair stored for the preceding unitig-occurrence. Finally, selecting for the successor-orientation pair (*s, ω*) in stab[*j*] yields *q*.

**Fig. S1.**
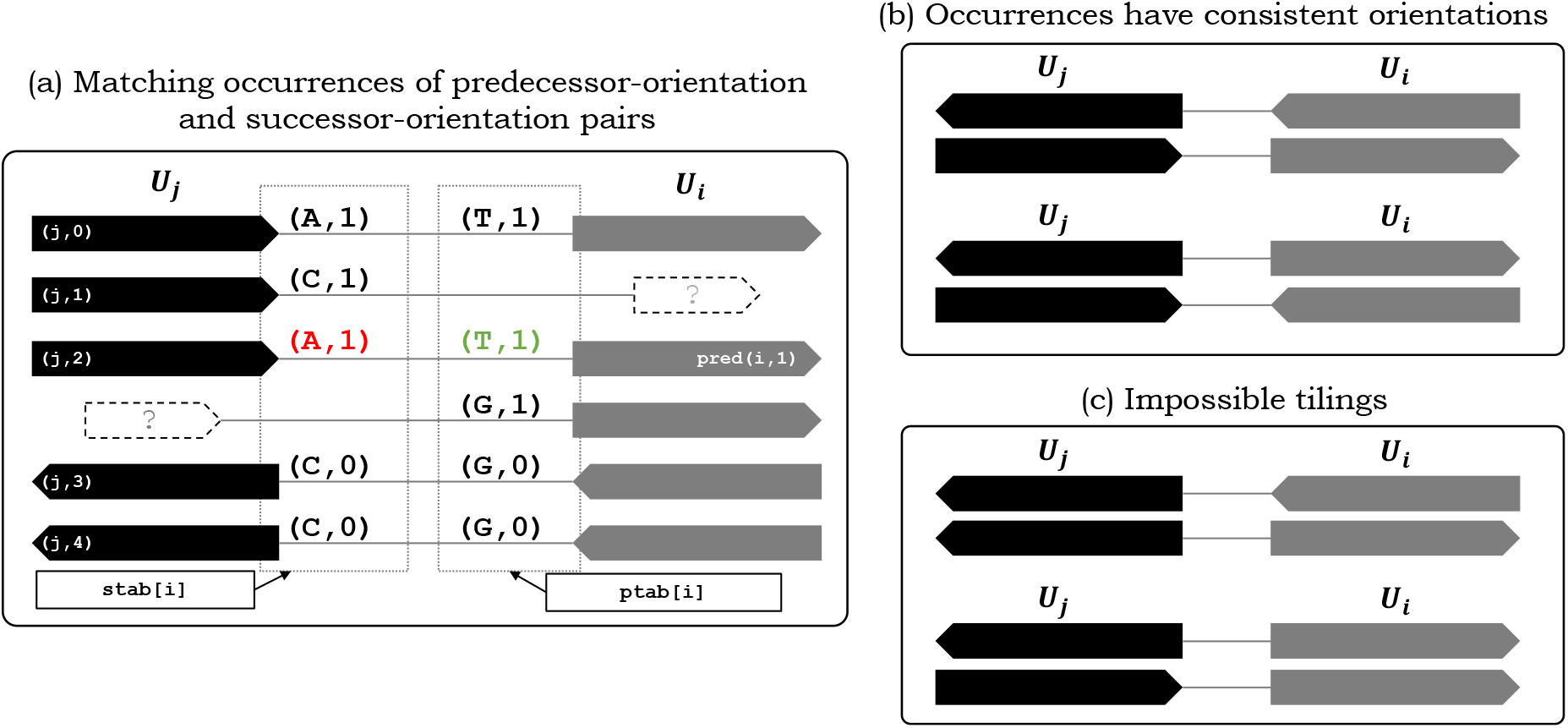
Properties of the `pred` query for unitig-tilings with orientations. (a) Adjacent pairs of successor and predecessor unitigs have consistent and unique co-occurring pairs of predecessor nucleotide-orientation successor nucleotide-orientation pairs. (b) Whenever a pair of unitigs occur adjacently on the tiling, the orientation of *one* fixes the orientation of the other (for odd *k*-mer sizes). (c) That is, if *U*_*j*_ with orientation *ω* precedes a unitig *U*_*i*_ with fixed orientation *o* once, it cannot precede another occurrence (of *U*_*i*_ with orientation *o*) in the opposite orientation.

##### Algorithm 5

The pred query with orientations

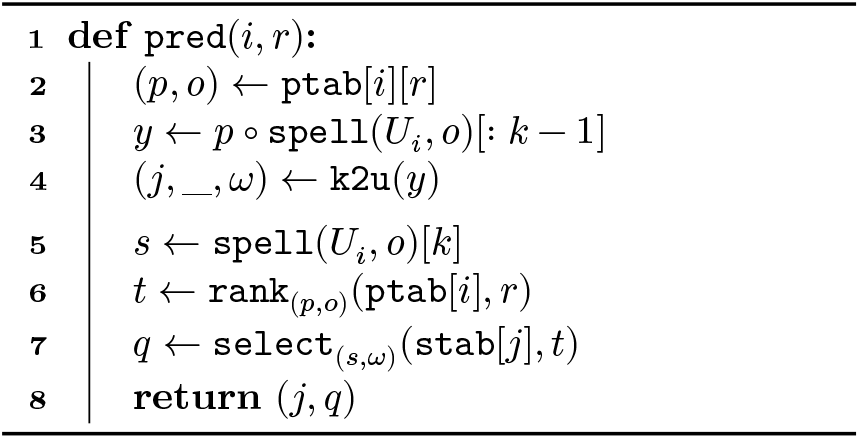

#### S.3.4 Unitig-unitig occurrences have consistent orientations and predecessor-successor nucleotides

The key to the correctness of `pufferfish2’s` reference tiling traversal, by way of successor-orientation and predecessor-orientation pairs, is that predecessor-successor nucleotide pairs for adjacent unitig-occurrences are consistent and unique up to orientation. Whenever unitigs *U*_*a*_ and *U*_*b*_ overlap and tile with some given *fixed* orientations, corresponding successor and predecessor nucleotides are consistent and always the same. Below, we prove Theorem 1 that formally states this property.

##### Theorem 1.

*Let unitigs U*_*a*_ *and U*_*b*_ *overlap and tile in orientations o and ω, with successor and predecessor nucleotides p and s. If any occurrence of U*_*a*_ *with orientation o is preceded by the nucleotide p, it must always be preceded by the same unitig U*_*b*_ *in the same orientation ω. Simultaneously, if any unitig U*_*b*_ *with orientation ω is succeeded by the nucleotide s, it must always be succeeded by the same unitig U*_*a*_ *in the same orientation o*.

*Proof*. Theorem 1 *is result of the lemmas proved below. Lemmas 1 and 2 state that with fixed orientations and predecessor and successor nucleotides, the identities of successor-predecessor unitig pairs must be unique. Lemmas 3 and 4 state that with fixed predecessor and successor nucleotides for fixed unitig identities, the orientations of a successor-predecessor unitig pair must be unique*.

##### Lemma 1

*Consider unitigs U*_*i*_, *U*_*j*_, *U*_*k*_ ∈ *𝒰. Let adjacent unitig occurrences T*_*a,b*_ = (*U*_*i*_, *o*) *and T*_*a,b*+1_ = (*U*_*j*_, *ω*) *occur with successor nucleotide s. For any c, d, there does not exist another pair of adjacent occurrences T*_*c,d*_ = (*U*_*i*_, *o*) *and T*_*c,d*+1_ = (*U*_*k*_, *ω*^′^) *with the same succeeding nucleotide s but with U*_*j*_ ≠ *U*_*k*_.

*Proof*. Let us assume the contrary. Let *z* be the last (*k* − 1)-mer on `spell`(*T*_*a,b*_), which is the same as `spell`(*T*_*c,d*_). Then the *k*-mer *z* ∘ *s* occurs on different unitigs *U*_*j*_ and *U*_*k*_. However, this is a contradiction since any unique *k*-mer occurs in only one unique unitig.

##### Lemma 2.

*Consider unitigs U*_*i*_, *U*_*j*_, *U*_*k*_ ∈ *𝒰. Let the occurrences T*_*a,b*_ = (*U*_*i*_, *o*) *and T*_*a,b*−1_ = (*U*_*j*_, *ω*) *occur with preceding nucleotide p. There does not exist another pair T*_*c,d*_ = (*U*_*i*_, *o*), *T*_*c,d*−1_ = (*U*_*k*_, *ω*^′^) *in ℛ where U*_*j*_ ≠ *U*_*k*_, *with the same preceding nucleotide s*.

*Proof*. This is symmetrical to Lemma 1.

##### Lemma 3

*Let* {*U*_*i*_, *U*_*j*_} ∈ *𝒰 Given unitig occurrences T*_*a,b*_ = (*U*_*i*_, *o*) *and T*_*a,b*+1_ = (*U*_*j*_, 1) *that tile R*_*a*_ *with successor nucleotide s. There does not exist another pair T*_*c,d*_ = (*U, o*), *T*_*c,d*+1_ = (*U*_*j*_, 0) *in ℛ with the same successor nucleotide s*.

*Proof*. Let us assume the contrary. Let *z* be the last (*k* − 1)-mer on *T*_*a,b*_ and *T*_*c,d*_. Suppose *z* ∘ *s* is the first *k*-mer on *U*_*i*_. The tiling on *R*_*c*_ implies that *z* ∘ *s* is the first *k*-mer on *U* _*j*_ and that 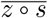 is the last *k*-mer on *Ū*_*j*_. But the tiling on *R*_*a*_ implies that *z* ∘ *s* is the first *k*-mer on *U*_*j*_. If |*U*_*j*_| = *k* and *U*_*j*_ is itself a *k*-mer, then the above implies *U*_*j*_ = *Ū*_*i*_. This cannot be the case, since we consider only odd-length *k*-mers, and no odd length *k*-mer can be equal to its reverse complement. If |*U*_*j*_| > *k*, then *z* occurs in two distinct positions in *U*_*j*_, this is again a contradication since any unique *k*-mer occurs in only one unique unitig.

##### Lemma 4

*Let* {*U*_*i*_, *U*_*j*_} ∈ *𝒰 Given unitig occurrences T*_*a,b*_ = (*U, o*) *and T*_*a,b*−1_ = (*V*, 1) *that tile R*_*a*_ *with predecessor nucleotide p. There does not exist another pair T*_*c,d*_ = (*U, o*), *T*_*c,d*−1_ = (*V*, 0) *in ℛ with the same precedecessor nucleotide p*.

*Proof*. This is symmetrical to Lemma 3.

### S.4 Constructing pufferfish2 from pufferfish

Building `pufferfish2` requires a linear scan over the *n* total unitig-occurrences in the tiling sequences indexed by a given `pufferfish` index to collect predecessor and successor nucleotides.The construction process is dominated by the time it takes to build the pair of wavelet matrices over all the predecessor and successor nucleotides of every unitig-occurrence. We note that since `pufferfish2` sparsifies and compresses and existing `pufferfish` index, it adopts `pufferfish’s` upstream preprocessing of unknown bases, where each N is replaced by a pseudo-random nucleotide. Constructing a wavelet matrix over an alphabet of size *σ* requires *𝒪*(*n* lg *σ*) time and amounts to successive stable partitions of the encoded characters according to their bitwise representations. In the future, we plan to update `pufferfish2` to index input reference sequences directly.

### S.5 Greedy unitig sampling with bounded traversal length *s*

Here we describe a *greedy* sampling scheme that greedily bounds traversal lengths to be at most of length *s*. To ensure that all backwards traversals terminate, the greedy sampling with integer paramater *s* first samples all unitigs that occur as the first occurrence of a tiling sequence, adds them to the set of sampled unitigs *𝒰*_*𝒮*_, and sets the corresponding bits in isSamp to 1 and all other bits to zero. Then, traversing tiling sequences in the order in which they appear, the greedy scheme maintains a counter of the distance to the last sampled unitig-occurrence. At each unitig occurrence *T*_*n,m*_ = *U*_*i*_, if *U*_*i*_ is already sampled (i.e., `isSamp`[*i*] is 1), the greedy scheme resets the counter to zero. Otherwise, the greedy scheme increases the counter by one. When the counter is greater than *s*, it samples the current unitig and resets the counter.

Although the greedy scheme is able to explicitly bound the traversal length it samples almost *all* unitigs when the length of tiling sequences become much larger than the number of unique unitigs. This is because, as implemented, `pufferfish2` samples *all occurrences* of a unitig, if said unitig is sampled. For example, when applying this sampling scheme to index a collection of seven human genomes, a greedy scheme with *s* = 3 samples 40% of unique unitigs that constitute more than 70% of unitig-*occurrences*. In this example, over 70% of `utab` must then be kept and uncompressed.

### S.6 Optimizations for pufferfish2

#### Caching traversals

When enumerating all positions of a unitig with `u2occ, pufferfish2` caches the `k2u` query — the empirically slowest constant-time operation in the `pred` query (Line 4 in Algorithm 3). The purpose of this k2u query is only to find the *identity* of the preceding unitig given a unitig-occurrence’s predecessor nucleotide. While a unitig may be preceded by *many* occurrences, preceding occurrences can have *at most* four unique unitig identities — one for each possible nucleotide. If *U*_*i*_ occurs more than once with *U*_*j*_ preceding it, *U*_*j*_ must always precede *U*_*i*_ with the same fixed predecessor nucleotide each time. For MRP queries, `pufferfish2` can avoid executing the redundant `k2u` queries (within `pred` queries) when the *same* nucleotide is prepended to different occurrences of *U*_*i*_. Specifically, during MRP queries where the `pred` query is executed, `pufferfish2` caches the mapping from predecessor nucleotides to preceding unitig identities. In practice, `pufferfish2` maintains an efficient LRU cache to memoize Lines 3 and 4 in Algorithm 3.

#### Caching streaming MRP queries

In practice, a *stream* of successive MRP queries for different *k*-mers often land in the same unitig (e.g. when querying *k*-mers on a sequenced read). So, instead of performing redundant `u2occ` queries that may perform backwards traversals for the same unitig, `pufferfish2` maintains a cache for the `u2occ` query. When successive *k*-mers are found to be in the same unitig via the `k2u` query, `pufferfish2` checks a “streaming cache” to avoid performing repeated `u2occ` queries for the same unitig. This caching scheme for “streaming” queries is also employed in [22].

#### Exiting early

In practice, programs such as read-mappers and aligners can exit early from the mapped reference position query if a queried *k*-mer is uninformative and occurs too frequently. With `pufferfish2`, instead of always computing the `u2occ` query for every occurrence in loop starting on Line 8, a caller of the MRP query can exit before the loop and avoid traversals altogether.

#### Interpolating between wavelet matrices and short linear scans

In practice, computing rank and select for short predecessor and successor nucleotide sequences (see Lines 6 and 7) is faster with a linear scan in an array than an operation in the wavelet matrix. So, for unitigs that occur at most 64 times, `pufferfish2` stores corresponding lists of nucleotides in packed arrays instead of wavelet matrices.

### S.7 Cost estimate for Amazon Web Services (AWS) EC2 instances

Estimated prices for AWS EC2 instances in the “US East” region are obtained from https://calculator.aws/#/estimate. EC2 instances were specified with 500Gb of storage and 8 CPUs for 10 hrs per week of usage. Recommended EC2 `x2gd.4xlarge` instances have 258GiB of memory whereas x2gd.2xlarge instances have 128GiB of memory. As of writing, estimated cost per month for `x2gd.4xlarge` and `x2gd.2xlarge` are 651USD and 351USD, respectively.

### S.8 Future work

1. Given the SPT definition, we have introduced strategies for sampling *𝒰*. That is, either a tile has *all* or *none* of its occurrences sampled. Yet, nothing theoretically prevents one from instead sampling over *𝒯*, so that *occurrences* are sampled according to their position on a tiling regardless of their unitig-identities. This approach introduces some extra complications but provides the benefit of allowing sampling schemes to *trivially* bound the worst-case traversal length, while also directly controlling the fraction of sampled entries by sampling every *s*-th occurrence. The question of what sampling strategy works better in practice is an interesting open question.
2. Does a smaller SPSS imply a smaller SPT? Currently, this is not clear, since working with unitigs dispenses entirely the space required for *𝒲, 𝔏*, and *𝒮*, so that a smaller SPSS may increase the space for representing the tiling given the need to encode *𝒲, 𝔏*, and *𝒮*.
3. We have provided an intuitive notion of how a “good” or “desirable” SPT looks: Ideally, an SPT amenable to indexing has few but long tiles and short tilings. Yet, rather than separating the problem of finding a set of tiles and then efficiently representing the tiling it induces, there is a more general optimization problem: Given a set *ℛ* of references, what is the SPT that minimizes the overall space, or the query time? Likewise, we may ask, if one has knowledge of the queries that are to be performed, how might the selection of samples be optimized to minimize the expected query time?
4. We have implemented one, specific, sparsification and compression scheme to reduce the size of SPTs. However, as hybrid encoding strategies have proven successful in optimizing the representation of *k*-mer-to-tile mappings [22], we may expect the same to be true of the tile-to-occurrence map. For example, long occurrence lists may compress well with traditional information retrieval compression schemes [31], and delta-encoding-like schemes may prove very effective in compressing the occurrence lists for tiles that almost always co-occur. In general, hybrid encoding and compression schemes likely hold great promise in tackling this problem.
5. Finally, we have considered here only exact and lossless representation of SPTs. However, many successful indexing schemes for problems like read mapping avoid indexing all sub-words, instead, for example, indexing only minimizers [32] or altering the sampling strategy in highly-repetitive regions. Thus, for many important applications it may not be necessary to have a *complete* and *lossless* index over the underlying SPT and it is possible that a *lossy* index over an SPT could be made much smaller and faster still.

## References

1. Fatemeh Almodaresi, Mohsen Zakeri, and Rob Patro. PuffAligner: a fast, efficient and accurate aligner based on the pufferfish index. Bioinformatics, June 2021. btab408.

2. Rob Patro, Geet Duggal, Michael I Love, Rafael A Irizarry, and Carl Kingsford. Salmon provides fast and bias-aware quantification of transcript expression. Nature Methods, 14(4):417–419, 2017.

3. Nicolas L Bray, Harold Pimentel, Páll Melsted, and Lior Pachter. Near-optimal probabilistic RNA-seq quantification. Nature Biotechnology, 34(5):525–527, 2016.

4. Rob Patro, Stephen M. Mount, and Carl Kingsford. Sailfish enables alignment-free isoform quantifi-cation from rna-seq reads using lightweight algorithms. Nature Biotechnology, 32(5):462–464, May 2014.

5. Travis Gagie, Gonzalo Navarro, and Nicola Prezza. Optimal-time text indexing in bwt-runs bounded space. In Proceedings of the Twenty-Ninth Annual ACM-SIAM Symposium on Discrete Algorithms, SODA ’18, page 1459–1477, USA, 2018. Society for Industrial and Applied Mathematics.

6. Massimiliano Rossi, Marco Oliva, Ben Langmead, Travis Gagie, and Christina Boucher. Moni: A pangenomic index for finding maximal exact matches. Journal of Computational Biology, 29(2):169– 187, 2022. PMID: 35041495.

7. Omar Ahmed, Massimiliano Rossi, Travis Gagie, Christina Boucher, and Ben Langmead. Spumoni 2: Improved pangenome classification using a compressed index of minimizer digests. bioRxiv, 2022.

8. Michael Burrows and David Wheeler. A block-sorting lossless data compression algorithm. In Digital SRC Research Report. Citeseer, 1994.

9. Fatemeh Almodaresi, Hirak Sarkar, Avi Srivastava, and Rob Patro. A space and time-efficient index for the compacted colored de bruijn graph. Bioinformatics, 34(13):i169–i177, 2018.

10. Daehwan Kim, Joseph M. Paggi, Chanhee Park, Christopher Bennett, and Steven L. Salzberg. Graph-based genome alignment and genotyping with hisat2 and hisat-genotype. Nature Biotechnology, 37(8):907–915, Aug 2019.

11. Erik Garrison, Jouni Sirén, Adam M. Novak, Glenn Hickey, Jordan M. Eizenga, Eric T. Dawson, William Jones, Shilpa Garg, Charles Markello, Michael F. Lin, Benedict Paten, and Richard Durbin. Variation graph toolkit improves read mapping by representing genetic variation in the reference. Nature Biotechnology, 36(9):875–879, Oct 2018.

12. Ilia Minkin, Son Pham, and Paul Medvedev. TwoPaCo: an efficient algorithm to build the compacted de Bruijn graph from many complete genomes. Bioinformatics, 33(24):4024–4032, 09 2016.

13. Rayan Chikhi, Antoine Limasset, and Paul Medvedev. Compacting de Bruijn graphs from sequencing data quickly and in low memory. Bioinformatics, 32(12):i201–i208, 06 2016.

14. Jamshed Khan and Rob Patro. Cuttlefish: fast, parallel and low-memory compaction of de Bruijn graphs from large-scale genome collections. Bioinformatics, 37(Supplement_1):i177–i186, 07 2021.

15. Jamshed Khan, Marek Kokot, Sebastian Deorowicz, and Rob Patro. Scalable, ultra-fast, and low-memory construction of compacted de bruijn graphs with cuttlefish 2. Genome Biology, 23(1):190, Sep 2022.

16. Barış Ekim, Bonnie Berger, and Rayan Chikhi. Minimizer-space de bruijn graphs: Whole-genome assembly of long reads in minutes on a personal computer. Cell Systems, 12(10):958–968.e6, 2021.

17. Mikhail Karasikov, Harun Mustafa, Gunnar Rätsch, and André Kahles. Lossless indexing with counting de bruijn graphs. Genome Res., 32(9):1754–1764, May 2022.

18. Amatur Rahman and Paul Medvedev. Representation of k-mer sets using spectrum-preserving string sets. In Russell Schwartz, editor, Research in Computational Molecular Biology, pages 152–168, Cham, 2020. Springer International Publishing.

19. Sebastian Schmidt and Jarno N. Alanko. Eulertigs: minimum plain text representation of k-mer sets without repetitions in linear time. bioRxiv, 2022.

20. Karel Břinda, Michael Baym, and Gregory Kucherov. Simplitigs as an efficient and scalable representation of de Bruijn graphs. Genome Biology, 22(1):96, April 2021.

21. Giulio Ermanno Pibiri. On weighted k-mer dictionaries. In International Workshop on Algorithms in Bioinformatics (WABI), pages 9:1–9:20, 2022.

22. Giulio Ermanno Pibiri. Sparse and skew hashing of k-mers. Bioinformatics, 38(Supplement_1):i185–i194, 06 2022.

23. Jarno N. Alanko, Simon J. Puglisi, and Jaakko Vuohtoniemi. Succinct k-mer sets using subset rank queries on the spectral burrows-wheeler transform. bioRxiv, 2022.

24. Francisco Claude and Gonzalo Navarro. The wavelet matrix. In International Symposium on String Processing and Information Retrieval, pages 167–179. onSpringer, 2012.

25. Alexander Bowe, Taku Onodera, Kunihiko Sadakane, and Tetsuo Shibuya. Succinct de Bruijn graphs. In International Workshop on Algorithms in Bioinformatics (WABI), pages o225–235. Springer, 2012.

26. Giulio Ermanno Pibiri and Rossano Venturini. Techniques for inverted index compression. ACM Comput. Surv., 53(6):125:1–125:36, 2021.

27. Uwe Baier, Timo Beller, and Enno Ohlebusch. Graphical pan-genome analysis with compressed suffix trees and the Burrows–Wheeler transform. Bioinformatics, 32(4):497–504, October 2015.

28. Pranvera Hiseni, Knut Rudi, Robert C. Wilson, Finn Terje Hegge, and Lars Snipen. HumGut: a comprehensive human gut prokaryotic genomes collection filtered by metagenome data. Microbiome, 9(1):165, July 2021.

29. Justin M. Zook, David Catoe, Jennifer McDaniel, Lindsay Vang, Noah Spies, Arend Sidow, Ziming Weng, Yuling Liu, Christopher E. Mason, Noah Alexander, Elizabeth Henaff, Alexa B.R. McIntyre, Dhruva Chandramohan, Feng Chen, Erich Jaeger, Ali Moshrefi, Khoa Pham, William Stedman, Tiffany Liang, Michael Saghbini, Zeljko Dzakula, Alex Hastie, Han Cao, Gintaras Deikus, Eric Schadt, Robert Sebra, Ali Bashir, Rebecca M. Truty, Christopher C. Chang, Natali Gulbahce, Keyan Zhao, Srinka Ghosh, Fiona Hyland, Yutao Fu, Mark Chaisson, Chunlin Xiao, Jonathan Trow, Stephen T. Sherry, Alexander W. Zaranek, Madeleine Ball, Jason Bobe, Preston Estep, George M. Church, Patrick Marks, Sofia Kyriazopoulou-Panagiotopoulou, Grace X.Y. Zheng, Michael Schnall-Levin, Heather S. Ordonez, Patrice A. Mudivarti, Kristina Giorda, Ying Sheng, Karoline Bjarnesdatter Rypdal, and Marc Salit. Extensive sequencing of seven human genomes to characterize benchmark reference materials. Scientific Data, 3(1):160025, June 2016.

30. Joan Mas-Lloret, Mireia Obón-Santacana, Gemma Ibáñez-Sanz, Elisabet Guinó, Miguel L. Pato, Francisco Rodriguez-Moranta, Alfredo Mata, Ana García-Rodríguez, Victor Moreno, and Ville Nikolai Pimenoff. Gut microbiome diversity detected by high-coverage 16S and shotgun sequencing of paired stool and colon sample. Scientific Data, 7(1):92, March 2020.

31. Alistair Moffat and Lang Stuiver. Binary interpolative coding for effective index compression. Information Retrieval, 3(1):25–47, 2000.

32. Heng Li. Minimap2: pairwise alignment for nucleotide sequences. Bioinformatics, 34(18):3094–3100, 2018.

