## Supplementary Materials for "Spectrum preserving tilings enable sparse and modular reference indexing"

### S.1 The mapped position query (MRP) for unitig-tilings

**Algorithm 4:** The MRP query for unitig-tilings

**def** `mrp`($x$):
    $tup \leftarrow$ `k2u`($x$)
    **if** $tup = \emptyset$ **then**
        **return** `[ ]`
    $(i, p) \leftarrow tup$
    $occ_i \leftarrow$ `num-occs`($U_i$)
    $ans \leftarrow$ `[ ]`
    **for** $r \leftarrow 0$ **to** $occs_i$ **do**
        $(n, s) \leftarrow$ `u2occ`($i, r$)
        $ans[r] = (n, s + p)$
    **return** ans

### S.2 Spectrum preserving tilings with *orientations*

We extend the definition of spectrum preserving tilings (without orientations) given in Section 3.1, to formally define spectrum preserving tilings (SPT) *with orientations*. An SPT with orientation allows tiles (members of a spectrum preserving string set) to occur in either a *forward* orientation as stored in memory as a nucleotide sequence, or a *backwards* orientation as the *reverse complement* of the stored sequence.

With respect to representing reference genomic sequences, using SPTs with orientations is particularly useful because it avoids redundantly encoding and storing occurrences of a $k$-mer *and* the reverse complement of said $k$-mer. Furthermore, since most sequencing technologies are agnostic to strands of DNA sequences, considering orientations enables the simultaneous and canonical representation of both corresponding strands of an indexed genomic sequence.

Also, as in [9], we consider only *odd* $k$-mer sizes so that no $k$-mer is its own reverse complement.

**Tiling sequences of tile *and orientation* pairs.** Given a fixed $k$-mer size, $k$, a tiling sequence $T_n$ in $\mathcal{T}$ is instead sequences of *tile-orientation* pairs where each occurrence is defined to be $T_{n,m} = (U_i, o)$, for some unitig $U_i \in \mathcal{U}$ and an orientation $o \in \{0, 1\}$. Here, $o = 1$ indicates that the unitig $U_i$ occurs in a forward orientation and $o = 0$ indicates that it occurs in the *backwards* orientation with reverse complement sequence $\overline{U_i}$. Notationally, $\overline{U_i}$ is the string that is the reverse complement of $U_i$ where $U_i$ is reversed and each nucleotide is replaced with its complement.

Let us define the $\texttt{spell}(U_i, o)$ function for a unitig orientation pair to return the forward sequence $U_i$ if $o = 1$ and the backwards, reverse complement sequence $\overline{U_i}$ otherwise. Abusing some notation, when $T_{n,m} = (U_i, o)$, let $\texttt{spell}(T_{n,m}) = \texttt{spell}(U_i, o)$.

Then formally, a spectrum preserving tiling with orientations for a reference collection $\mathcal{R} = \{R_1, \ldots, R_N\}$ tiles each reference sequence $R_n$ with sequences *spelled by* occurrences of tile-orienation pairs. Specifically, each $R_n$ can be reconstructed by gluing together the sequences that tile-orientation pairs *spell*. For each $R_n$ that is tiled by $M_n$ tile occurrences, the SPT satisfies the property that:

$$R_n = \texttt{spell}(T_{n,1})[w_{n,1} : w_{n,1} + l_{n,1}] \;\oplus_k\; \ldots \;\oplus_k \texttt{spell}(T_{n,M_n})[w_{n,M_n} : w_{n,M_n} + l_{n,M_n}]$$

#### S.2.1 Returning queries with orientations

Accordingly, when indexing an SPT with orientations the mapped reference position query, $k$-mer-to-tile query, and tile-to-occurrence query also return orientations. Here, we extend and reintroduce the queries defined in Section 3.

1. **The mapped reference position (MRP) query** Given any $k$-mer $x$, the MRP query enumerates the positions and orientations of all occurrences of $x$ in $\mathcal{R}$. Precisely, each returned occurrence is a tuple $(n, p, o)$, that specifies that $k$-mer $x$ occurs in reference $n$ at position $p$ with orientation $o$. That is, if $o = 1$, then $x$ occurs in the forward orientation as $R_n[p : p+k] = x$. Otherwise, the reverse complement occurs as $R_n[p : p+k] = \overline{x}$. If a $k$-mer does not occur in some $R_n \in \mathcal{R}$, the query returns an empty list.
2. **The kmer-to-tile query**: Given a $k$-mer $x$, $\texttt{k2tile}(x)$ returns $(i, p, o)$ — the identity of the tile $U_i$ that contains $x$, the offset (position) into the tile $U_i$ where $x$ occurs, and the *orientation* of how $x$ occurs. That is, $\texttt{k2tile}(x) = (i, p, 1)$ if $U_i[p : p+k] = x$, and $\texttt{k2tile}(x) = (i, p, 0)$ if $U_i[p : p+k] = \overline{x}$ where $x$ occurs in the backwards orientation as the reverse complement. If $x$ is not in $\mathcal{R}$, `k2tile`, $\texttt{k2tile}(x)$ returns $\emptyset$.
3. **The tile-to-occurrence query**: Given the $r$-th occurrence of the tile $U_i$, $\texttt{tile2occ}(i, r)$ returns the tuple $(n, o, s, w, l)$ that encodes how *and in what orientation* $U_i$ tiles the reference $R_n$. Let the $r$-th occurrence of $U_i$ be a tile-occurrence $T_{n,m}$ on $\mathcal{T}$ where $T_{n,m} = U_i, o$ for some orientation $o$. Then $\texttt{tile2occ}(i, r)$ returns $(n, o, s_{n,m}, w_{n,m}, l_{n,m})$. When $\texttt{tile2occ}(i, r) = (n, o, s, w, l)$ and $o = 1$, the $r$-th occurrence of $U_i$ occurs on $R_n$ at position $(s + w)$, with the sequence $U_i[w : w + l]$. When $\texttt{tile2occ}(i, r) = (n, o, s, w, l)$ and $o = 0$, the $r$-th occurrence of $U_i$ occurs on $R_n$ at position $(s + w)$, with the sequence $\overline{U_i}[w : w + l]$.

With some arithmetic bookkeeping considering orientations and lengths, the MRP query with orientations can again be decomposed into the two corresponding $k$-mer-to-tile and tile-to-occurrence queries that also return orientations. Although not introduced with respect to an SPT, the `pufferfish` index developed by Almodaresi et al. [9] is implemented exactly this way as an index over an SPT *with orientations* of unitigs.

### S.3 `Pufferfish2`: the `pred` query with orientations

`Pufferfish2`'s sampling scheme and the `pred` query can be applied when considering orientations — our implemented tool does exactly this. Below, we extend Section 4 to fully specify the introduced sampling scheme and the `pred` query when orientations are considered.

#### S.3.1 Predecessor and successor nucleotides

When SPT references, predecessor and successor nucleotides are defined and obtained with respect to sequences on the *references*. Specifically, the predecessor nucleotide is the first nucleotide of the last $k$-mer on of the preceding unitig-occurrence *as spelled with the corresponding orientation* of the occurrence. The successor nucleotide is defined in the same manner.

Suppose $T_{n,m} = (U_i, o)$, and $T_{n,m-1} = (U_j, \omega)$, and let the unitigs have lengths $\ell_i$ and $\ell_j$, respectively. We say that, $T_{n,m-1}$ precedes $T_{n,m}$ with predecessor nucleotide $p$ *and orientation* $o$. Concretely, $p$ is the first nucleotide on the last $k$-mer of the preceding unitig, with $p = \texttt{spell}(T_{n,m-1})[\ell_j - k]$. We say that, $T_{n,m}$ succeeds $T_{n,m-1}$ with successor nucleotide $s$ *and orientation* $\omega$. Accordingly, the successor nucleotide, $s$, is the last nucleotide on the first $k$-mer of the succeeding unitig, with $s = \texttt{spell}(T_{n,m})[k]$.

#### S.3.2 Storing nucleotide-orientation pairs in ptab and stab

Instead of storing only nucleotides, `pufferfish2` stores nucleotide-orientation pairs in implementation. That is, for each occurrence $T_{n,m} = (U_i, o)$ that is the $r$-th occurrence of a not-sampled unitig $U_i$,

$$\texttt{ptab}[i][r] = (\texttt{pred-nuc}(T_{n,m}), o).$$

And for each occurrence $T_{n,m} = (U_i, o)$ that is the $r$-th occurrence of *any* unitig $U_i$,

$$\texttt{stab}[i][r] = (\texttt{succ-nuc}(T_{n,m}), o).$$

In summary, `ptab` and `stab` store for each corresponding unitig-occurrence, the nucleotides that succeed and precede it as they occur on a tiled reference, *and* the orientation of said occurrence.

#### S.3.3 Computing the pred query by matching ranks of predecessor-orientation and successor-orientation pairs

When orientations are considered, computing the `pred` query requires matching ranks of predecessor-orientation and successor-orientation pairs. Critically, any time a pair of unitigs occur as a successor-predecessor pair in *fixed* orientations, the corresponding pair of predecessor and successor nucleotides are *consistent* and also *fixed*. Furthermore, if an occurrence of the $U_j$ in orientation $\omega$ precedes a unitig $U_i$ with orientation $o$, *any other* occurrence of $U_j$ that precedes $U_i$ with orientation $o$ must also occur with orientation $\omega$. We state and prove this property with Theorem 1 and illustrate examples of both possible and impossible unitig-occurrences with Fig. S1.

Theorem 1 guarantees that whenever $U_i$ occurs with orientation $o$ with predecessor nucleotide $p$ preceded by $U_j$, $U_j$ must occur with fixed orientation $\omega$ with a fixed successor nucleotide $s$. We illustrate this correspondence in Fig. S1. Algorithm 5 implements `pred` with orientations considered.

To find the identity, $j$, of the preceding unitig occurrence, Algorithm 5 must construct the last $k$-mer of the corresponding occurrence $U_j$ as it appears on the reference. Specifically, in Line 3 it spells $U_i$ before extracting the overlapping $(k-1)$-mer. Here, `k2u` returns orientation, $\omega$, of the queried $k$-mer on $U_j$, which must also be the orientation of the preceding unitig occurrence on the reference. Furthermore, the successor nucleotide, $s$, for the preceding occurrence of $U_j$ must be the $k$-th nucleotide on $U_i$ spelled with orientation $o$ (Line 5).

Now, Algorithm 5 has all it needs to compute $q$, the unitig-rank of the preceding occurrence of $U_j$. Computing the rank of $(p, o)$ in $\texttt{ptab}[i]$ yields the rank of the corresponding successor-orientation

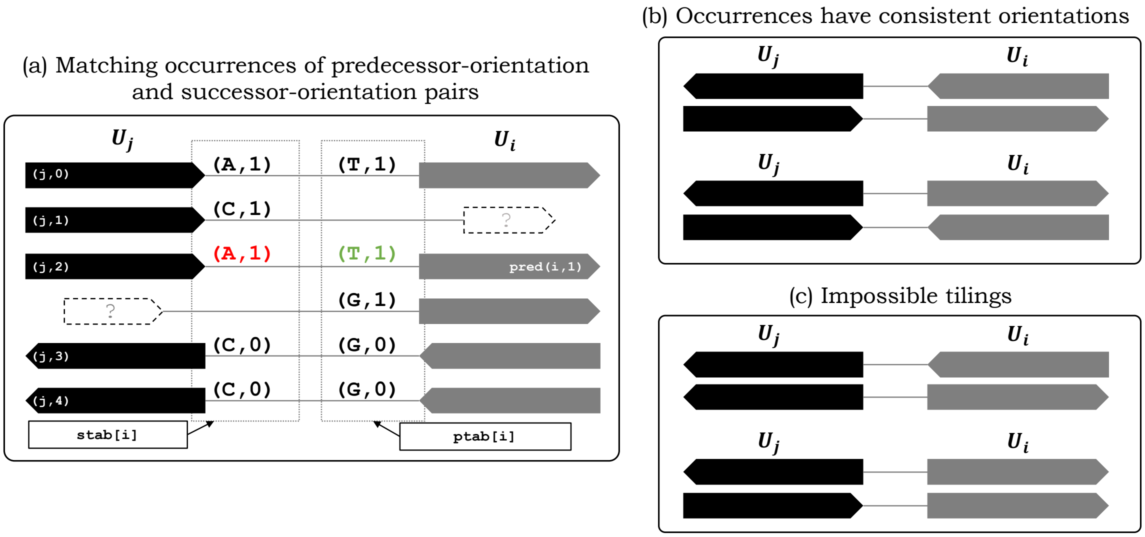


**Fig. S1.** Properties of the `pred` query for unitig-tilings with orientations. (a) Adjacent pairs of successor and predecessor unitigs have consistent and unique co-occurring pairs of predecessor nucleotide-orientation successor nucleotide-orientation pairs. (b) Whenever a pair of unitigs occur adjacently on the tiling, the orientation of *one* fixes the orientation of the other (for odd $k$-mer sizes). (c) That is, if $U_j$ with orientation $\omega$ precedes a unitig $U_i$ with fixed orientation $o$ once, it cannot precede another occurrence (of $U_i$ with orientation $o$) in the opposite orientation.

pair stored for the preceding unitig-occurrence. Finally, selecting for the successor-orientation pair $(s, \omega)$ in `stab`$[j]$ yields $q$.

**Algorithm 5:** The `pred` query with orientations

**def** `pred`$(i, r)$**:**
- $(p, o) \leftarrow$ `ptab`$[i][r]$
- $y \leftarrow p \circ$ `spell`$(U_i, o)[: k-1]$
- $(j, \_, \omega) \leftarrow$ `k2u`$(y)$
- $s \leftarrow$ `spell`$(U_i, o)[k]$
- $t \leftarrow$ `rank`$_{(p,o)}$(`ptab`$[i], r)$
- $q \leftarrow$ `select`$_{(s,\omega)}$(`stab`$[j], t)$
- **return** $(j, q)$

#### S.3.4 Unitig-unitig occurrences have consistent orientations and predecessor-successor nucleotides

The key to the correctness of `pufferfish2`'s reference tiling traversal, by way of successor-orientation and predecessor-orientation pairs, is that predecessor-successor nucleotide pairs for adjacent unitig-occurrences are consistent and unique up to orientation. Whenever unitigs $U_a$ and $U_b$ overlap and tile with some given *fixed* orientations, corresponding successor and predecessor nucleotides are consistent and always the same. Below, we prove Theorem 1 that formally states this property.

**Theorem 1.** *Let unitigs $U_a$ and $U_b$ overlap and tile in orientations $o$ and $\omega$, with successor and predecessor nucleotides $p$ and $s$. If any occurrence of $U_a$ with orientation $o$ is preceded by the nucleotide $p$, it must always be preceded by the same unitig $U_b$ in the same orientation $\omega$. Simultaneously, if any unitig $U_b$ with orientation $\omega$ is succeeded by the nucleotide $s$, it must always be succeeded by the same unitig $U_a$ in the same orientation $o$.*

*Proof.* Theorem 1 is result of the lemmas proved below. Lemmas 1 and 2 state that with fixed orientations and predecessor and successor nucleotides, the identities of successor-predecessor unitig pairs must be unique. Lemmas 3 and 4 state that with fixed predecessor and successor nucleotides for fixed unitig identities, the orientations of a successor-predecessor unitig pair must be unique.

**Lemma 1.** *Consider unitigs $U_i, U_j, U_k \in \mathcal{U}$. Let adjacent unitig occurrences $T_{a,b} = (U_i, o)$ and $T_{a,b+1} = (U_j, \omega)$ occur with successor nucleotide $s$. For any $c, d$, there does not exist another pair of adjacent occurrences $T_{c,d} = (U_i, o)$ and $T_{c,d+1} = (U_k, \omega')$ with the same succeeding nucleotide $s$ but with $U_j \neq U_k$.*

*Proof.* Let us assume the contrary. Let $z$ be the last $(k-1)$-mer on $\texttt{spell}(T_{a,b})$, which is the same as $\texttt{spell}(T_{c,d})$. Then the $k$-mer $z \circ s$ occurs on different unitigs $U_j$ and $U_k$. However, this is a contradiction since any unique $k$-mer occurs in only one unique unitig.

**Lemma 2.** *Consider unitigs $U_i, U_j, U_k \in \mathcal{U}$. Let the occurrences $T_{a,b} = (U_i, o)$ and $T_{a,b-1} = (U_j, \omega)$ occur with preceding nucleotide $p$. There does not exist another pair $T_{c,d} = (U_i, o)$, $T_{c,d-1} = (U_k, \omega')$ in $\mathcal{R}$ where $U_j \neq U_k$, with the same preceding nucleotide $s$.*

*Proof.* This is symmetrical to Lemma 1.

**Lemma 3.** *Let $\{U_i, U_j\} \in \mathcal{U}$ Given unitig occurrences $T_{a,b} = (U_i, o)$ and $T_{a,b+1} = (U_j, 1)$ that tile $R_a$ with successor nucleotide $s$. There does not exist another pair $T_{c,d} = (U, o)$, $T_{c,d+1} = (U_j, 0)$ in $\mathcal{R}$ with the same successor nucleotide $s$.*

*Proof.* Let us assume the contrary. Let $z$ be the last $(k-1)$-mer on $T_{a,b}$ and $T_{c,d}$. Suppose $z \circ s$ is the first $k$-mer on $U_i$. The tiling on $R_c$ implies that $\overline{z \circ s}$ is the first $k$-mer on $\overline{U_j}$ and that $z \circ s$ is the last $k$-mer on $U_j$. But the tiling on $R_a$ implies that $z \circ s$ is the first $k$-mer on $U_j$. If $|U_j| = k$ and $U_j$ is itself a $k$-mer, then the above implies $U_j = \overline{U_i}$. This cannot be the case, since we consider only odd-length $k$-mers, and no odd length $k$-mer can be equal to its reverse complement. If $|U_j| > k$, then $z$ occurs in two distinct positions in $U_j$, this is again a contradication since any unique $k$-mer occurs in only one unique unitig.

**Lemma 4.** *Let $\{U_i, U_j\} \in \mathcal{U}$ Given unitig occurrences $T_{a,b} = (U, o)$ and $T_{a,b-1} = (V, 1)$ that tile $R_a$ with predecessor nucleotide $p$. There does not exist another pair $T_{c,d} = (U, o)$, $T_{c,d-1} = (V, 0)$ in $\mathcal{R}$ with the same precededecessor nucleotide $p$.*
